# The vascular niche controls *Drosophila* hematopoiesis via Fibroblast Growth Factor signaling

**DOI:** 10.1101/2020.11.26.400127

**Authors:** Manon Destalminil-Letourneau, Ismaël Morin-Poulard, Yushun Tian, Nathalie Vanzo, Michèle Crozatier

## Abstract

In adult mammals, hematopoiesis, the production of blood cells from hematopoietic stem and progenitor cells (HSPCs), is tightly regulated by extrinsic signals from the microenvironment called “niche”. Bone marrow HSPCs are heterogeneous and controlled by both endosteal and vascular niches. The *Drosophila* hematopoietic lymph gland is located along the cardiac tube which corresponds to the vascular system. In the lymph gland, the niche called Posterior Signaling Center controls only a subset of the heterogeneous hematopoietic progenitor population indicating that additional signals are necessary. Here we report that the vascular system acts as a second niche to control lymph gland homeostasis. The FGF ligand Branchless produced by vascular cells activates the FGF pathway in hematopoietic progenitors. By regulating intracellular calcium levels, FGF signaling maintains progenitor pools and prevents blood cell differentiation. This study reveals that two niches contribute to the control *of Drosophila* blood cell homeostasis through their differential regulation of progenitors.

## Introduction

In adult mammals, HSPCs in the bone marrow ensure the constant renewal of blood cells. The cellular microenvironment of HSPCs, called “niche”, regulates hematopoiesis under both homeostatic and immune stress conditions (Asada et al., 2017; Calvi et al., 2003; Calvi and Link, 2015; He et al., 2014; Kiel et al., 2005; Kobayashi et al., 2016; Morrison and Scadden, 2014; Zhao and Baltimore, 2015). Recent studies have revealed significant molecular and functional heterogeneity within the HSPC pool (for review (Haas et al., 2018)). These findings challenge the differential contribution of niche cell types to HSPC diversity. Given the high conservation of regulatory networks between insects and vertebrates, *Drosophila* has become an important model to study how hematopoiesis is controlled (Evans et al., 2003; Hartenstein, 2006). Insect blood cells, or hemocytes, are related to the mammalian myeloid lineage. In *Drosophila,* three blood cell types are produced: plasmatocytes that are macrophages involved in phagocytosis, crystal cells involved in melanisation and wound healing and lamellocytes required for encapsulation of pathogens too large to be destroyed by phagocytosis. Lamellocytes represent a cryptic cell fate since they only differentiate at the larval stage and in response to specific immune challenges such as wasp parasitism (Lemaitre and Hoffmann, 2007). The lymph gland is the larval hematopoietic organ and is composed of paired lobes, one pair of anterior lobes and several pairs of posterior lobes, aligned along the anterior part of the cardiac tube (CT) which corresponds to the vascular system (Fig. 1a and (Lanot et al., 2001)). In third instar larvae the anterior lobes comprise three zones: a medullary zone (MZ) containing hematopoietic progenitors, a cortical zone (CZ) composed of differentiated blood cells, and a small group of cells called the Posterior Signaling Center (PSC) (Fig. 1a and (Crozatier et al., 2004; Jung et al., 2005)). The PSC produces a variety of signals that regulate lymph gland homeostasis (for review see (Banerjee et al., 2019; Letourneau et al., 2016; Yu et al., 2018)). Recently we established that cardiac cells produce the ligand Slit which, through the activation of Robo receptors in the PSC, controls the proliferation and clustering of PSC cells and in turn their function (Morin-Poulard et al., 2016). Furthermore, the MZ progenitor population is heterogeneous and a subset of progenitors called “core progenitors” which express the *knot/Collier (Kn/Col*) and the *thioester-containing protein-4* (*tep4*) genes is aligned along the cardiac tube and are maintained independently from the PSC (Fig. 1a and (Baldeosingh et al., 2018; Benmimoun et al., 2015; Oyallon et al., 2016)). Altogether, these data led us to ask whether signals derived from cardiac cells are involved in the control of lymph gland homeostasis, i.e. the balance between progenitors and differentiated blood cells, independently from the PSC. To address this possibility we performed a candidate RNAi screen in cardiac cells to identify new potential signaling pathways involved in the crosstalk between the vascular and the hematopoietic organs.

**Figure 1:**
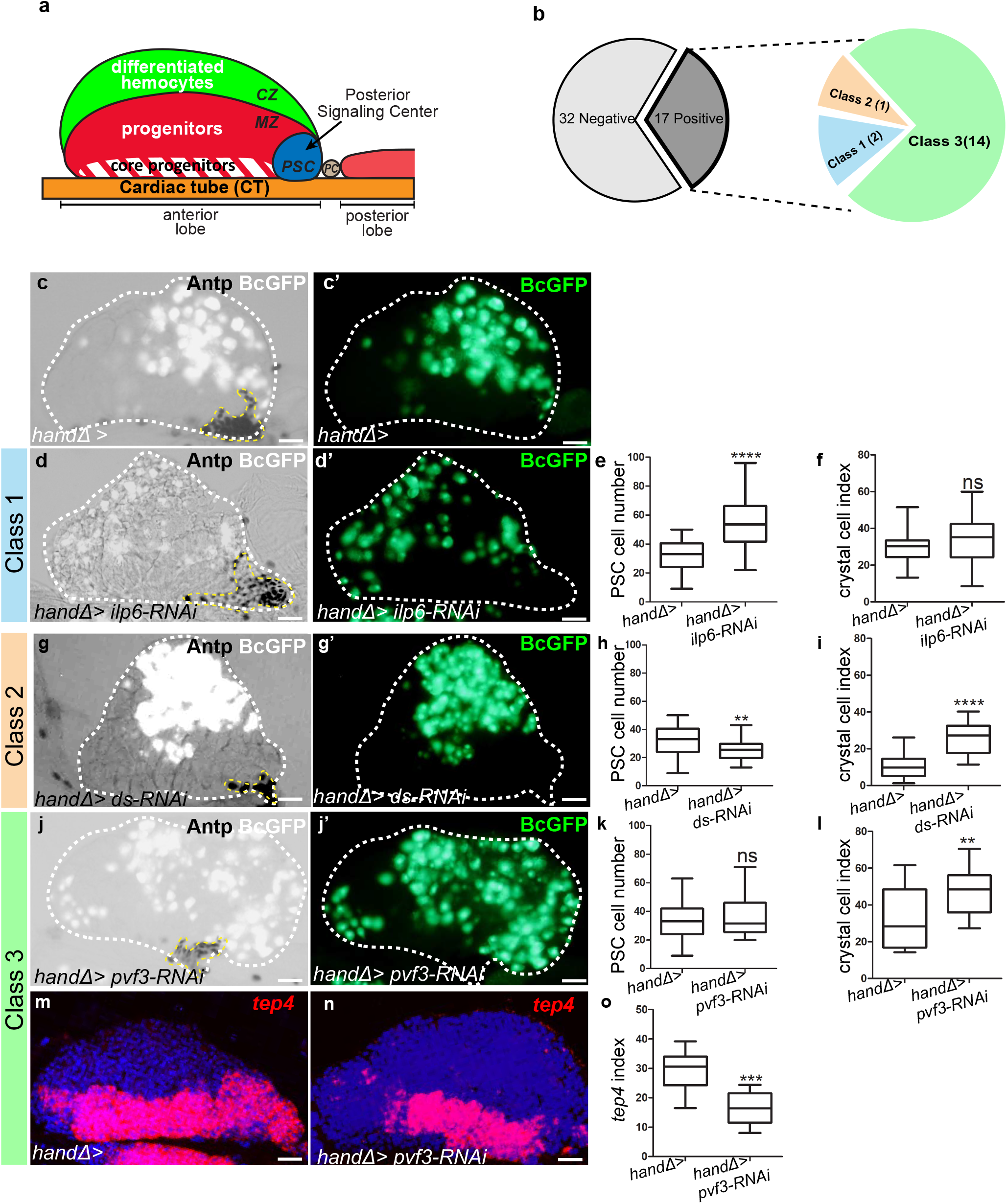
Lymph gland organization and RNAi screen results. (a) Representation of lymph gland anterior and posterior lobes from third instar larvae. The anterior lobe is composed of progenitors (red) and core progenitors (hatched red), and the cortical zone (CZ, green). The PSC is blue and the cardiac tube (CT) /vascular system, is orange. PC corresponds to pericardial cell. (b) Summary of the screen performed by expressing RNAi in cardiac cells using the *handΔ-gal4* driver. The number of genes corresponding to the different classes of phenotype is given. Subsequent panels illustrate the control and observed lymph gland defects (c, d, g, j). Anterior lobe and PSC are delimited by white and yellow dashed lines, respectively. Black cell-GFP (BcGFP, white) labels crystal cells and Antp (black) the PSC. (c’, d’, g’, j’) BcGFP is in green; (e, h, k) PSC cell numbers; (f, i, l) Crystal cell index. (c-f) Reducing *ilp6* in cardiac cells (d, d’) augments PSC cell number (e) without affecting crystal cell differentiation (f); this defines class 1. (g-i) Knocking down *dachsous (ds)* in cardiac cells (g, g’) decreases PSC cell number (h) and increases crystal cell index (i); this defines class 2. (j-l) Reducing *pvf3* in cardiac cells (j, j’) does not modify PSC cell number (k) but increases crystal cell differentiation (l); this defines class 3. (m, n) *tep4* (red) labels core progenitors. Decrease in *tep4* expression is observed when *pvf3* is knocked down in cardiac cells. (o) *tep4* index. For all quantifications and figures, statistical analysis *t*-test (Mann-Whitney nonparametric test) was performed using GraphPad Prism 5 software. Error bars represent SEM and *P<0,1;** P< 0,01; ***P <0,001; ****P <0,0001 and ns (not significant). In all confocal pictures nuclei are labelled with Topro (blue) and scale bars = 20μm.

Here we show that several signals produced by cardiac cells contribute to maintain lymph gland homeostasis. We investigated in more detail the role of the Fibroblast Growth Factor (FGF) ligand Branchless (Bnl). FGF signaling is conserved during evolution and is less complex in *Drosophila* than in humans. Ligand binding to a FGF receptor (FGFR) promotes its dimerization, which results in its tyrosine-phosphorylation, thus providing a scaffold to recruit different partners (Muha and Muller, 2013; Ornitz and Itoh, 2015). In mammals, ligand binding to the FGFR activates Ras/Raf-Mek-MAPK, PI3K/AKT and PLCγ-Ca^2+^ signaling pathways (Turner and Grose, 2010).The *Drosophila* genome encodes two FGF receptors, Breathless (Btl) and Heartless (Htl), and three ligands, Bnl, Thisbe (Ths) and Pyramus (Pyr), (Beiman et al., 1996; Glazer and Shilo, 1991; Gryzik and Muller, 2004; Klambt et al., 1992; Sutherland et al., 1996). Htl is activated by Ths and Pyr, while Btl is activated by Bnl. We established that Bnl is expressed in cardiac cells and signals to its receptor Breathless (Btl) expressed in progenitors. Bnl/Btl-FGF activation controls progenitor intracellular Ca^2+^ concentration, probably by activating Phospholipase Cγ (PLCγ) which regulates endoplasmic reticulum Ca^2+^ stores. Altogether, these data strongly support the conclusion that the cardiac tube plays a role similar to a niche by regulating lymph gland hematopoiesis.

## Results

### A cardiac screen identifies genes controlling lymph gland homeostasis

To investigate the role of cardiac cells in the control of lymph gland hematopoiesis, we performed a functional screen based on the expression, in cardiac cells, of RNAis directed against transcripts encoding known *Drosophila* ligands. For this we used the cardiac *handΔ-gal4* driver which is expressed in cardiac cells throughout the three larval stages (Figure 1-figure supplement 1a-c’ and (Monier et al., 2005; Morin-Poulard et al., 2016)) to screen a collection of RNAi lines corresponding to 49 *Drosophila* ligands (Figure 1-figure supplement 2). As read-outs, we analyzed blood cell differentiation with the crystal cell reporter BcGFP (Tokusumi et al., 2009), and PSC cell numbers and morphology by performing Antennapedia (Antp) immunostaining (Mandal et al., 2007). Compared to the control, 17 RNAi lines showed lymph gland homeostasis defects that were classified into three groups (Fig. 1b). Class 1: Increased PSC cell numbers but no effect on crystal cell differentiation (Fig. 1c-f). Two RNAis against *ilp6* and *spätzle4* transcripts belong to this class (Figure 1-figure supplement 2). Class 2: Decreased PSC cell numbers and increased crystal cell differentiation (Fig. 1g-i). Only one RNAi against *dachsous* (*ds*) belongs to this class (Figure 1-figure supplement 2). Class 3: No effect on PSC cell numbers but increased crystal cell differentiation (Fig. 1j-l); 14 RNAis belong to this class. The class 3 phenotype strongly suggested that signals from cardiac cells could control crystal cell differentiation independently from the PSC. We extended the analysis of the 14 corresponding genes by labeling the core progenitors with *tep4 in situ* hybridization (Krzemien et al., 2007). Reduced *tep4* expression was observed for 12 RNAi treatments out of 14 (Fig. 1m-o and Figure 1-figure supplement 2), indicating that the corresponding genes are required in cardiac cells to maintain *tep4* expression in lymph gland progenitors and to prevent crystal cell differentiation. To avoid any bias due to the *handΔ-gal4* driver, we also tested *NP1029-gal4*, an independent cardiac cell driver (Figure 1-figure supplement 1d-f’ and (Monier et al., 2005; Morin-Poulard et al., 2016)). Among the 14 RNAi candidates, 9 gave a similar phenotype with both drivers (Figure 1-figure supplement 2). In conclusion, our functional screen allowed us to identify 9 ligands involved in communication between cardiac cells and hematopoietic progenitors to control lymph gland homeostasis.

### The FGF ligand Bnl from cardiac cells controls lymph gland homeostasis

One candidate identified in our screen was Bnl. Previous studies have shown that Htl-FGF signaling is required during both early embryogenesis for lymph gland specification (Grigorian et al., 2011; Mandal et al., 2004) and in L3 larvae to control lymph gland progenitors (Dragojlovic-Munther and Martinez-Agosto, 2013). However, no role for *bnl* in the lymph gland has been described yet. Since *bnl* knock-down in cardiac cells significantly enhanced crystal cell differentiation in the lymph gland we decided to pursue an analysis of the Bnl-FGF pathway. Since Bnl is a diffusible ligand, we first documented *bnl* mRNA expression by *in situ* hybridization. *bnl* is expressed in cardiac and pericardial cells (Fig. 2 a-a’’), in agreement with previously published data (Jarecki et al., 1999). We also observed a weak *bnl* expression in MZ progenitors (as labelled by domeMESO>GFP in Figure 2 a-a”), in differentiating hemocytes (as labelled by hml>GFP) and in a subset of crystal cells (marked by BcGFP) whereas no expression was detected in the PSC (Figure 2-figure supplement 1a-c”). In a heterozygous *bnl* loss-of-function mutant context where one copy of *bnl (bnl^P2^/+)* is missing, we observed an increased number of crystal cells compared to the control (Fig. 2b-d). To specifically knock down *bnl* in cardiac cells we expressed *bnl-RNAi* under the control of the cardiac tube specific driver *handΔgal4*. *bnl* loss-of-function experiments were performed from the L2 larval stage on, after the cardiac tube had formed, to avoid possible cardiac tube morphological defects (see MM and Figure 2-figure supplement 1d-e). *bnl* down-regulation in cardiac cells resulted in increased differentiation of both crystal cells and plasmatocytes (Fig. 2e-f, i and Figure 2-figure supplement 1f-h). Increased crystal cell differentiation was also observed using another independent *bnl* RNAi line (Figure 2-figure supplement 1i-k) and with the alternative *NP1029-gal4* driver (Figure 2-figure supplement 1l-n). Applying *bnl* knockdown only after the L2 stage by using the GAL80 ^ts^ system (McGuire et al., 2004) led to a similar crystal cell differentiation defect (Figure 2-figure supplement 1 o-q). We then analyzed MZ progenitors when *bnl* was knocked down in cardiac cells, using DomeMESO-RFP that labels all progenitors, and *tep4* and Col that are expressed in the core progenitors (Krzemien et al., 2007; Oyallon et al., 2016). Compared to wild-type, a reduced expression of the three markers was observed in *handΔ>bnl-RNAi* lymph glands (Fig. 2j-r), indicating that Bnl from cardiac cells non-cell autonomously controls MZ progenitor maintenance. Altogether, these data indicate that Bnl produced in the cardiac tube acts in third instar larvae to control lymph gland homeostasis.

**Figure 2:**
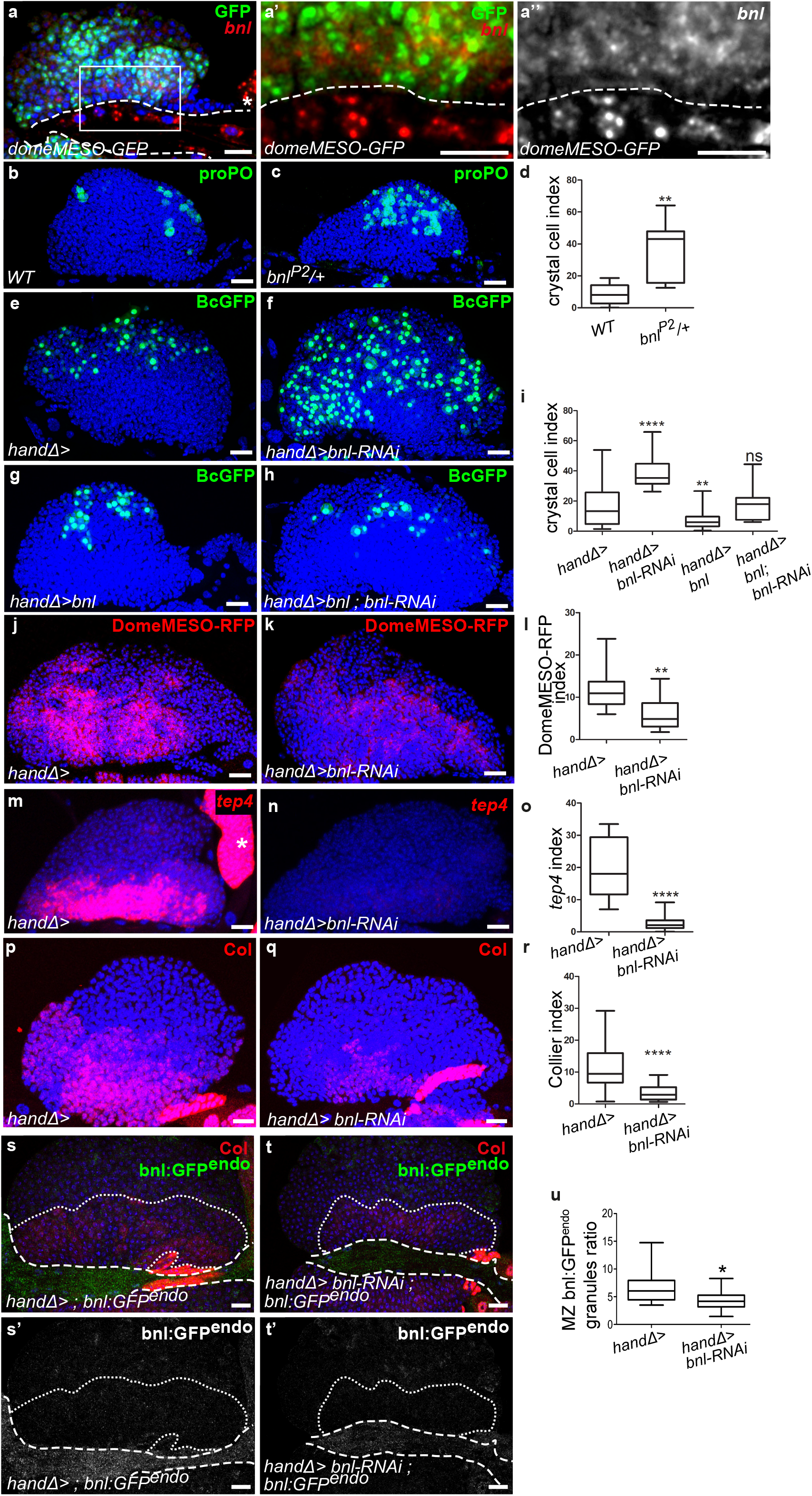
Ligand Bnl is expressed in cardiac cells and controls lymph gland homeostasis. (a) A maximum projection of 5 confocal lymph gland sections, *bnl* (red) is expressed in cardiac cells and MZ progenitors that express domeMESO-GFP (green). (a’, a’’) An enlarged view, *bnl* is red (a’) or white (a”). A white dashed line indicates the cardiac tube. * indicates a pericardiac cell. (b, c) proPO (green) labels crystal cells. *bnl^P2^/+* heterozygous mutant lymph glands have an increased number of crystal cells (c) compared to the control (b). (e-f, g-h) Black-cell GFP (BcGFP, green) labels crystal cells. (d, i) Crystal cell index. Co-expression of *bnl* and *bnl-RNAi* in cardiac cells restores the wildtype number of crystal cells (i). (j, k) DomeMESO-RFP (red) labels MZ progenitors. Compared to the control (j) barely detectable DomeMESO-RFP levels are observed when *bnl* is knocked down in cardiac cells (k). (l) DomeMESO-RFP index. (m, n) *tep4* labels core progenitors. Compared to the control (m) lower levels of *tep4* (red) are observed when *bnl* is knocked down in cardiac cells (n). (o) *tep4* index. (p-q) Col labels core progenitors. Compared to the control (p) lower levels of Col are observed in the core progenitors when *bnl* is knocked down in cardiac cells (q). (r) Col index. (s-t’) Maximum projection of 5 confocal sections of the lymph gland expressing *bnl:GFP ^endo^* (green) and Col immunostaining that labels MZ progenitors (red). Compared to the control (s, s’) a decrease in bnl:GFP ^endo^ in green (t) and white (t’) is observed when *bnl* is knocked down in cardiac cells. Fine and thick dashed lines indicate the MZ and CT contours, respectively. (u) Bnl:GFP^endo^ granules ratio in the MZ.

Since *bnl* is transcribed in MZ progenitors, though at low levels, we also analyzed its function in these cells. Reduction of *bnl* expression in progenitors (*dome>bnl-RNAi)* led to a significant increase in crystal cell differentiation as well as a decrease in *tep4* expression (Figure 2-figure supplement 2a-f), indicating that Bnl produced by MZ progenitors is required to maintain their identity and to prevent their differentiation. Altogether, these data show that Bnl is produced by both MZ and cardiac cells and that both sources are required in the control of lymph gland homeostasis.

To determine whether Bnl produced by cardiac cells contributes to the pool of Bnl present in the MZ, we analyzed endogenous Bnl distribution. We used the bnl:GFP^endo^ knock-in allele that recapitulates *bnl* expression (Du et al., 2018). In agreement with *in situ bnl* detection, bnl:GFP^endo^ was found in cardiac cells and in MZ progenitors (Figure 2s-s’). However, when *bnl* was knocked-down only in the cardiac tube (*handΔ>bnl RNAi, bnl:GFP^endo^*; Figure 2 t-u) we overserved a reduction of *bnl:GFP^endo^* both in cardiac cells and MZ progenitors thus establishing that Bnl produced by cardiac cells contributes to the global MZ Bnl pool. The concomitant increased hemocyte differentiation suggests that Bnl levels contributed by the heart are required for lymph gland homoeostasis. To further support this conclusion, we overexpressed *bnl* only in cardiac cells (*handΔ>bnl)* which led to reduced crystal cell numbers (Fig 2i), and this confirms that the level of Bnl produced by cardiac cells controls hemocyte differentiation. Finally, rescue of crystal cell numbers (Fig. 2g-i) and of progenitor marker expression (Figure 2-figure supplement 2g-i) was observed with a simultaneous expression of *bnl* and *bnl-RNAi* in cardiac cells (*handΔ>bnl; bnl-RNAi*). Altogether, these data establish that Bnl produced by cardiac cells is required for lymph gland hematopoiesis.

Since the PSC controls lymph gland cell differentiation (Benmimoun et al., 2015; Morin-Poulard et al., 2016; Oyallon et al., 2016; Tokusumi et al., 2010), we also looked at PSC cells by analyzing the expression of the PSC marker Antp (Mandal et al., 2007) when *bnl* was downregulated in cardiac cells. No PSC cell number or clustering defects were observed (Figure 2-figure supplement 2j-l). Hh expression in the PSC regulates progenitors and blood cell differentiation (Baldeosingh et al., 2018; Mandal et al., 2007; Tokusumi et al., 2010). The *hhF4-GFP* reporter transgene (Tokusumi et al., 2010) was expressed in PSC cells in the *handΔ>bnl-RNAi* context similar to the control (Figure 2-figure supplement 2m-o). These data strongly suggest that cardiac cell Bnl neither affects PSC cell numbers nor Hh activity, and likely acts directly on MZ progenitors to control lymph gland homeostasis. Altogether, these data indicate that although it is transcribed in many lymph gland cells, *bnl* expression in cardiac cells plays an essential role in the control of lymph gland homeostasis.

### The FGF receptor Btl expressed in progenitors, controls lymph gland homeostasis

Bnl activates the FGF pathway by binding Btl (Kadam et al., 2009). To document endogenous Btl expression in the lymph gland, we used the btl:cherry^endo^ knock-in allele which recapitulates Btl expression (Du et al., 2018). Strong btl:cherry^endo^ expression was observed in cardiac cells and lower levels in MZ progenitors (as labelled by domeMESO-GFP in Fig 3a-a”). Whereas no expression was detected in PSC cells, a very faint expression occurs in a subset of crystal cells and in most differentiating blood cells (labelled by BcGFP and Hml>GFP, respectively, in Figure 3-figure supplement 1a-c”).

**Figure 3:**
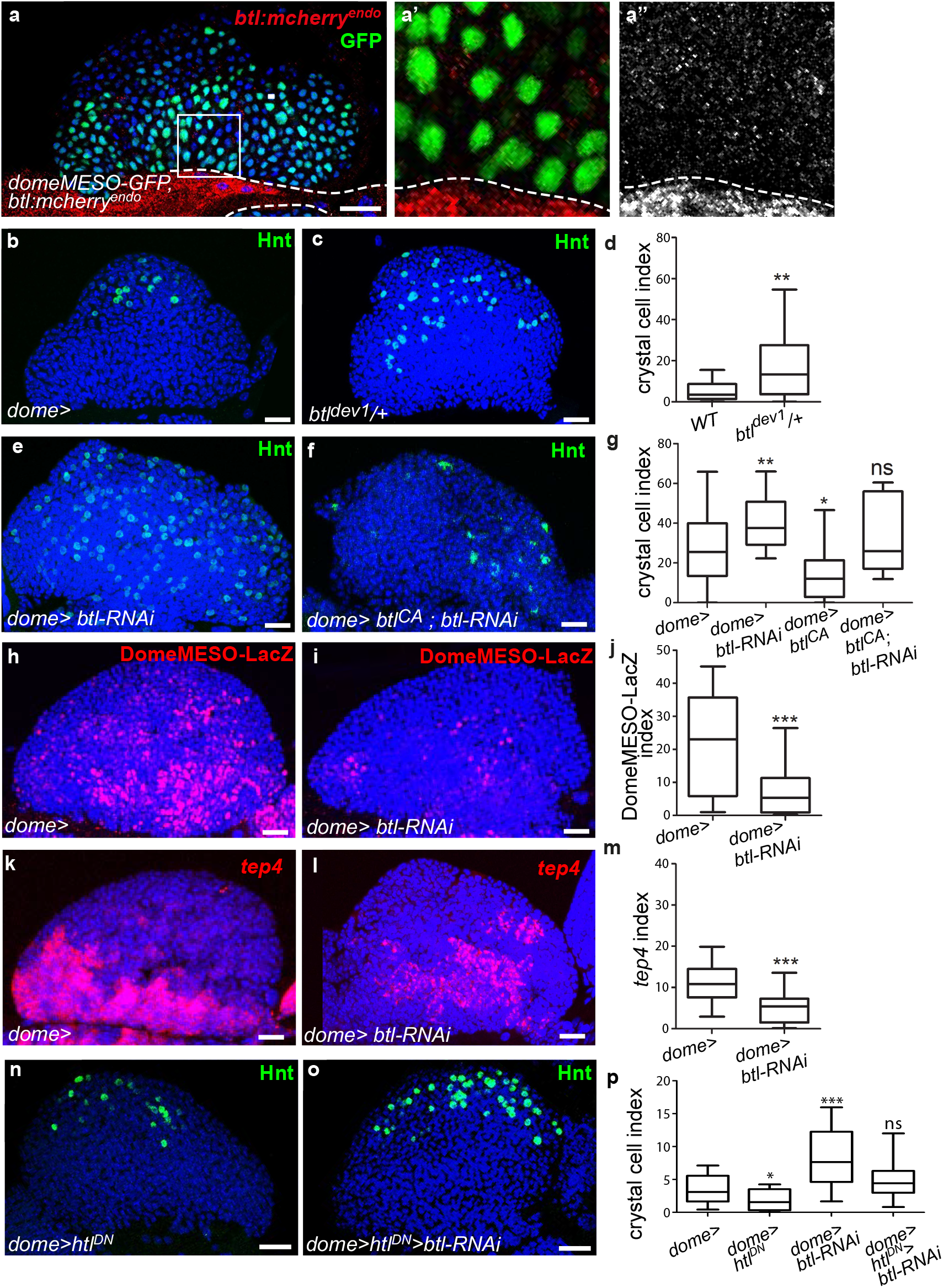
Receptor Btl is expressed in hematopoietic progenitors and required to control lymph gland homeostasis. (a) A maximum projection of 5 confocal lymph gland sections of larvae expressing *btl:cherry^endo^* (red) and *domeMESO-GFP* that labels MZ progenitors (green). (a’, a’’) An enlarged view, *btl:cherry^endo^* red (a’) or white (a”). Dashed lines indicate the cardiac tube contour. *btl:cherry^endo^* is expressed in cardiac cells and MZ progenitors. (b-c, e-f) Hindsight (Hnt, green) labels crystal cells. Crystal cell differentiation is increased in *btl^dev1^/+* heterozygous mutant larvae (c) compared to the control (b). (e, f) Crystal cell numbers increase when *btl* is knocked down in progenitors (e) and crystal cell differentiation is rescued when a constitutive activated *btl* receptor (*btl^CA^*) is expressed in the *btl-RNAi* context (f). (d, g) Crystal cell index. (h, i) DomeMESO-LacZ (red) labels MZ progenitors. Compared to the control (h) barely detectable domeMESO-LacZ levels are observed when *btl* is knocked down in progenitors (i). (k, l) Lower levels of *tep4* (red) are observed when *btl* is knocked down in progenitors (l) compared to the control (k). (j, m) DomeMESO-LacZ and tep4 index, respectively. (n, p) Crystal cell numbers decrease when a dominant negative *htl* receptor (*htl^DN^*) is knocked down in progenitors (n) and crystal cell differentiation is increased when *htl^DN^* is co-expressed with *btl-RNAi* (o). (p) Crystal cell index.

To determine whether *btl* is required for lymph gland homeostasis, we looked at crystal cell differentiation in a heterozygous loss-of-function mutant context, where one copy of *btl* is mutated (*btl^dev1^/+*). The resulting crystal cell index was higher than in controls (Fig. 3b-d), revealing that *btl* controls lymph gland homeostasis. We then knocked down *btl* by expressing RNAi in either MZ progenitors (*dome>)* or cardiac cells (*handΔ>). btl* downregulation in cardiac cells did not significantly affect crystal cell numbers or MZ progenitors (Figure 3 - figure supplement 1d-i). In contrast, knocking down *btl* in MZ progenitors led to increased crystal cell (Fig. 3e, g) and plasmatocyte numbers (Figure 3-figure supplement 1j-l), together with a reduced expression of the two progenitor markers *domeMESO-LacZ* and *tep4* (Fig. 3h-m). We then performed rescue experiments with a constitutively active form of Btl (*btl^CA^*, (Pares and Ricardo, 2016)). The expression of *btl^CA^* in progenitors (*dome>btl^CA^)* led to reduced crystal cell numbers, i.e., a phenotype opposite to that of *btl* loss-of-function (Fig. 3g). The co-expression of *btl^CA^* and *btl-RNAi* in progenitors (dome>*btl*-RNAi>*btl^CA^*; Fig. 3f, g) rescued crystal cell differentiation, confirming that Btl is required in MZ progenitors. Finally, we examined whether PSC cells were affected when *btl* was knocked down in progenitors. No difference in PSC cell numbers or clustering was observed compared to the control (Figure 3-figure supplement 1m-o). We conclude that *btl* expression in MZ progenitors is required to control lymph gland homeostasis.

As opposed to Btl-FGF inhibition, Htl-FGF pathway knock-down in MZ progenitors was reported to block blood cell differentiation (Dragojlovic-Munther and Martinez-Agosto, 2013). To investigate the relationship between the two pathways in the MZ, we performed epistasis experiments. Expression of a dominant–negative form of Htl (dome>Htl^DN^) in progenitors led to a decrease in crystal cell differentiation, in agreement with a previous report (Dragojlovic-Munther and Martinez-Agosto, 2013). Simultaneous expression of Htl^DN^ and *btl-RNAi* (*dome>Htl^DN^>btl-RNAi*) restored a wildtype number of crystal cells (Figure 3n-p). These data indicate that there is no hierarchy between Btl-FGF and Htl-FGF pathways. They also suggest that their simultaneous activity in MZ progenitors ensures a robust regulation of hemocyte differentiation.

In conclusion, downregulating Btl in MZ progenitors causes a defect in lymph gland homeostasis similar to that caused by Bnl downregulation in cardiac cells. This strongly suggests that Bnl/Btl-FGF signaling mediates inter-organ communication between MZ progenitors and the vascular system. This leads us to propose that by acting directly on MZ progenitors, the cardiac tube plays a role similar to a niche.

### Bnl secreted by cardiac cells is taken up by lymph gland progenitors

Bnl originating from cardiac cells and acting on MZ progenitors raised the question of its mode of diffusion. To investigate this question, we expressed a functional GFP-tagged version of Bnl (*UAS-Bnl::GFP,* (Lin and Affolter, 2009)) in cardiac cells. In addition to the expected GFP detection in these cells, discrete GFP positive cytoplasmic punctate dots/granules were detected in MZ progenitors (Fig. 4a-a’’), indicating that Bnl::GFP can propagate from cardiac to lymph gland cells. Many cytoplasmic Bnl-GFP positive punctate dots in the MZ were Btl:Cherry positive (Figure 4b-b”). To further characterize these Bnl-GFP dots, we labelled recycling vesicles and late endosomes, using the ubi-Rab11-cherryFP reporter and Rab7 immunostaining, respectively. We found that many Rab11-positive and Rab7-positive vesicles co-localized with Bnl-GFP in the MZ (Figure 4c-d”). The simplest explanation is that Bnl::GFP secreted by cardiac cells is internalized by MZ progenitors, likely through receptor mediated endocytosis.

**Figure 4:**
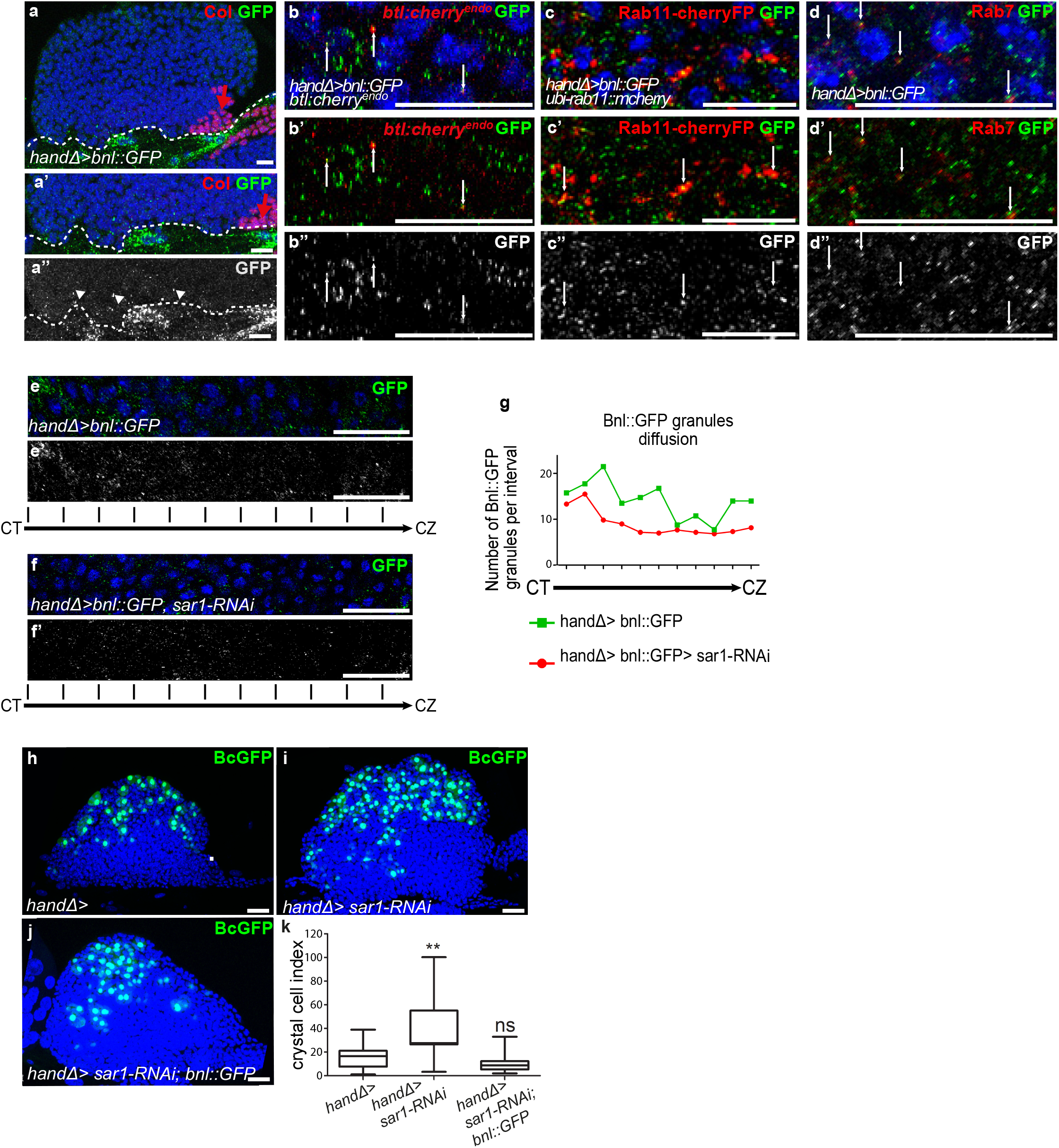
Ligand Bnl secreted by cardiac cells controls lymph gland crystal cell differentiation. (a) Active bnl::GFP fusion protein is expressed in cardiac cells using *handΔ-gal4* driver. Dashed lines indicate cardiac tube and the PSC is labelled by Collier (Col, red and red arrow). (a’, a’’) An enlarged view; Bnl::GFP is green (a’) or white (a”). Bnl::GFP positive granules are detected in cardiac and lymph gland cells (arrowheads). (b-b”) Enlargement of MZ area close to the cardiac tube in larvae expressing Bnl::GFP fusion protein (green) in cardiac cells (*handΔ-gal4>Bnl::GFP*) and Btl:mcherry^endo^ (red). Bnl::GFP cytoplasmic punctate dots (green in b-b’ and white in b”) co-localize with Btl:mcherry^endo^ (yellow and arrows in b’). (c,c”) Enlargement of MZ area close to the cardiac tube in larvae expressing *ubi-Rab11cherryFP* (red), a marker for recycling endocytic vesicles; Bnl::GFP fusion protein (green) is expressed in cardiac cells (*handΔ-gal4>Bnl::GFP*). Bnl::GFP cytoplasmic punctate dots (green in c-c’ and white in c”) co-localize with ubi-Rab11cherryFP (yellow and arrows in c’). (d-d”) Enlargement of MZ area close to the cardiac tube in larvae expressing Bnl::GFP fusion protein (green) in cardiac cells (*handΔ-gal4>Bnl::GFP*) and Rab7 immunostainings (red in d, d’ and white in d”). (d-d’) Bnl::GFP cytoplasmic punctuate dots co-localize with Rab7 positive dots (yellow and arrows in d’). (e-f’) Enlargement of lymph gland cross sections extending from the cardiac tube (CT) to the cortical zone (CZ). Bnl::GFP fusion protein, expressed in cardiac cells (*handΔ-gal4>Bnl::GFP*) is green (e, f) and white (e’, f’). Knocking down *sar1* in cardiac cells (f, f’) leads to a decrease in Bnl::GFP cytoplasmic punctate dots compared to the control (e, e’). (g) Quantification of Bnl::GFP cytoplasmic punctate dots/granules. (h, j) BcGFP (green) labels crystal cells. Knocking down *sar1* in cardiac cells (i) increases crystal cell numbers compared to the control (h). Crystal cell differentiation rescue is observed when bnl::GFP is co expressed with *sar1-RNAi* (j, k). (k) Crystal cell index.

To further confirm the role of Bnl secreted by cardiac cells, we impaired endoplasmic reticulum (ER) vesicle formation by knocking down the Secretion-associated Ras-related GTPase1 (Sar1) specifically in cardiac cells. The Sar1GTPase plays a key role in the biogenesis of transport vesicles and acts by regulating vesicular trafficking (Saito et al., 2017; Yorimitsu et al., 2014). The simultaneous expression of *sar1-RNAi* and Bnl::GFP in cardiac cells using the *handΔ-gal4* driver (*handΔ>bnl::GFP>sar1-RNAi*) resulted in reduced bnl::GFP cytoplasmic punctate dots in MZ progenitors compared to the control (Figure 4e-g), further confirming that cardiac cells secrete Bnl. Previous data had established that the Slit ligand, produced by cardiac cells, activates Robo receptors in the PSC and as a consequence controls PSC cell proliferation and clustering (Morin-Poulard et al., 2016). Consistent with the impairment of cardiac cell secretion when *sar*1 is knocked down in cardiac cells (*handΔ> and NP1029> sar1-RNAi*, Figure 4-figure supplement 1a-f), we also observed an increase in PSC cell numbers as well as a slight defect in their clustering, likely an effect of Slit/Robo impairment. These data indicate that *sar1* knock-down in cardiac cells impairs their secretory capacity and therefore Bnl secretion..

We then analyzed the consequences on lymph gland blood cell differentiation. When *sar1* was knocked down in cardiac cells using *handΔ* or NP1029 drivers (Fig. 4h-i, k and Figure 4 - figure supplement 1g-i) crystal cell numbers were higher than the control, indicating that cardiac cell secretion capacity is necessary for lymph gland hematopoiesis. To determine whether the overexpression of *bnl* in cardiac cells can compensate for decreased secretion, we performed rescue experiments. Crystal cell differentiation was improved when *sar1-RNAi* and bnl::GFP were simultaneously expressed in cardiac cells (Fig. 4j, k). In conclusion, these results indicate that Bnl secreted by cardiac cells is likely taken up by MZ progenitors to activate Btl-FGF signaling, which in turn regulates lymph gland homeostasis.

### FGF activation in progenitors regulates their calcium levels

The next step was to address how Bnl/Btl-FGF signaling in progenitors controls lymph gland homeostasis. Depending on the cellular context, the FGF pathway activates the MAPK or PI3K pathways, or PLCγ that controls intracellular Ca^2+^ levels (Ornitz and Itoh, 2015). Inactivation of MAPK or PI3K in lymph gland progenitors leads to a phenotype opposite to that of knock-down of *btl* in progenitors or of *bnl* in cardiac cells (Dragojlovic-Munther and Martinez-Agosto, 2013). This strongly suggests that Bnl/Btl-FGF signaling in progenitors does not involve MAPK or PI3K activity. It was previously reported that intracellular Ca^2+^ levels regulate hematopoietic progenitor maintenance: reduction of cytosolic Ca^2+^ in lymph gland progenitors leads to the loss of progenitor markers and to increased blood cell differentiation (Shim et al., 2013). Since Bnl/Btl-FGF knock-down and reduction of Ca^2+^ in progenitors induce similar lymph gland defects, we asked whether both mechanisms were functionally linked. We investigated Ca^2+^ levels within MZ progenitors using the Ca^2+^ sensor GCaMP3, which emits green fluorescence only with high Ca^2+^levels (Nakai et al., 2001). This sensor is expressed under the control of the *dome-gal4* driver. In agreement with previous reports, high Ca^2+^ levels were detected in MZ progenitors (Fig. 5a and (Shim et al., 2013)). Knocking down *btl* in MZ progenitors starting from L1 stage led in third instar larvae to decreased fluorescence compared to the control, indicating a reduction in Ca^2+^ levels (Fig. 5a-c). Since no difference with control larvae could be observed at L2 stage (Figure5-Figure Supplement1 a-c) we conclude that the Bnl/Btl-FGF pathway is not required for MZ progenitor specification but is required in third instar larvae to regulate Ca^2+^ levels. We then asked whether restoring high Ca^2+^ levels in progenitors could rescue the lymph gland defects due to reduced Bnl/Btl-FGF function. When free Ca^2+^ was high within the progenitors, such as following overexpression of Calmodulin dependent kinase II (CaMKII; *dome>CaMKII)* or of IP3R (Tep4>UAS-IP3R) which controls ER-mediated Ca^2+^ release into the cytosol (Shim et al., 2013), a significant decrease in crystal cell numbers was observed compared to the control (Fig. 5d-h and Figure 5-figure supplement 1d-h). This indicates that high Ca^2+^ levels in progenitors inhibit blood cell differentiation, which is in agreement with previously published data (Shim et al., 2013). Simultaneous *btl* reduction and Ca^2+^ increase in progenitors, through CaMKII or IP3R overexpression (*dome> CamKII*;*btl-RNAi* Fig. 5g-h and *tep4>IP3R>btl-RNAi;* Figure 5-figure supplement 1g-h), leads to a decrease in crystal cell numbers compared to the sole *btl-RNAi* knockdown. Overall, these data suggest that in MZ progenitors, regulation of Ca^2+^ levels functions downstream of the Bnl/Btl-FGF pathway. In vertebrates, FGF activation can recruit and activate PLCγ, which induces Ca^2+^ release from the ER (Ornitz and Itoh, 2015). *small wing* (*sl*) encodes the sole *Drosophila* PLCγ homologue (Thackeray et al., 1998). To investigate the role of PLCγ in the lymph gland, we analyzed crystal cell differentiation in strong hypomorphic *sl^2^* mutants. Compared to the control, increased crystal cell differentiation was observed (as labelled by Hnt (Fig. 5 j, m). We then addressed the role of *PLCγ* specifically in progenitors. Knocking down PLCγ in progenitors (*dome>sl-RNAi)* led to increased crystal cell differentiation compared to the control (Fig. 5i, l), revealing that PLCγ in MZ progenitors regulates lymph gland hemocyte differentiation. Finally, we performed epistasis experiments to decipher the relationship between *sl* and the Btl-FGF pathway. When *btl^CA^* was expressed in progenitors in a *sl^2^* mutant context (*sl^2^; dome>blt^CA^*; Figure 5 k, n), the crystal cell index was similar to that in the *sl^2^* mutant alone. These data establish that *sl* acts downstream of the Bnl/Btl-FGF pathway. Altogether, our data support the hypothesis that in MZ progenitors, Bnl/Btl-FGF signaling leads to the activation of PLCγ, which controls Ca^2+^ levels and in turn hemocyte differentiation.

**Figure 5:**
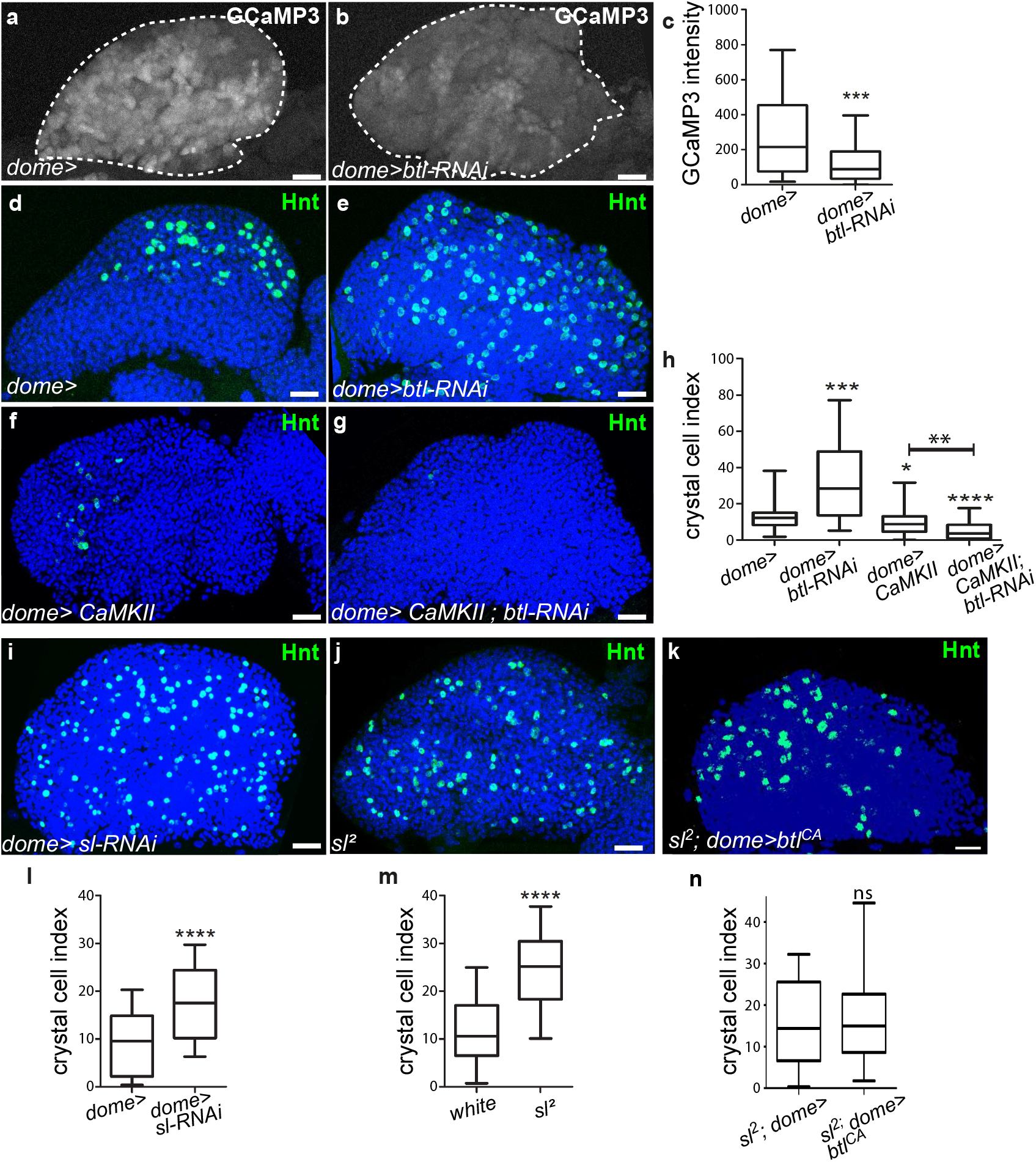
Btl receptor interacts genetically with CamKII to control blood cell differentiation by preventing high Ca^2+^ levels in progenitors. (a, b) GCaMP3 Ca^2+^ sensor (*dome>UAS-GCaMP3*) is white. GCaMP3 intensity decreases when *btl is* knocked down in MZ progenitors (b) compared to the control (a). (c) Quantification of GCaMP3 intensity. (d-g, i-k) Hnt (green) labels crystal cells. Crystal cell differentiation decrease is observed when Ca^2+^ levels increased due to CaMKII expression in progenitors (*dome>CaMKII,* f) compared to the control (d). Co-expression of CaMKII and *btl* RNAi in progenitors (*dome>CaMKII; btl-RNAi*, g) leads to a decrease in crystal cell number compared to the *btl* knock-down alone (e). (h, l-n) Crystal cell index. Crystal cell differentiation increase is observed in *sl^2^* homozygous mutant larvae (j, m) and when *sl* is knocked down in progenitors (i, l) compared to the control (d). (k, n) No difference in crystal cell index is observed in *sl^2^* homozygous mutant larvae and in a *sl^2^* homozygous mutant where btl^CA^ is expressed in MZ progenitors *(sl^2^; dome>btl^CA^*).

## Discussion

The control of HSPCs by a specific microenvironment called “niche” is established both in mammals and in *Drosophila*. The niche is defined by its capacity to directly regulate, through signals, stem cells and progenitors. In the mammalian bone marrow HSPCs are under the control of the endosteal and vascular niches (Asada et al., 2017; Calvi et al., 2003; Calvi and Link, 2015; He et al., 2014; Kiel et al., 2005; Morrison and Scadden, 2014). In *Drosophila,* lymph gland studies have so far concentrated on the PSC acting as a niche. However, a subset of lymph gland progenitors (core-progenitors), which express Col and *tep4* and are aligned along the cardiac tube, is maintained in the lymph gland even when the PSC function is impaired, suggesting that other signals alongside those from the PSC are required (Baldeosingh et al., 2018; Benmimoun et al., 2015; Oyallon et al., 2016). Here, we report that cardiac cells play a role similar to a niche, since they directly control core progenitor maintenance. We show that communication between the vascular system and the lymph gland involves Bnl/Btl-FGFsignaling. Bnl secreted by cardiac cells activates Bnl/Btl-FGF in progenitors, which in turn controls hemocyte homeostasis. Our data indicate that Bnl/Btl-FGF signaling regulates lymph gland homeostasis by controlling calcium levels in progenitors via PLCγ activation (Figure 6). In a previous study, we showed that signals from the cardiac tube, namely Slit, can act on the PSC, but that no cellular communication between the cardiac tube and MZ progenitors is involved (Morin-Poulard et al., 2016). Now we establish that cardiac cells regulate the extent of progenitor differentiation in the lymph gland. Therefore, two separate niches (the PSC and the cardiac tube) control lymph gland homeostasis. While the PSC acts only on a subset of MZ progenitors (Baldeosingh et al., 2018; Oyallon et al., 2016), the cardiac tube directly regulates core progenitors and in turn all MZ progenitors (Figure 6).The identification of two niches that differentially regulate lymph gland progenitors sheds further light on the parallels existing between *Drosophila* lymph gland and mammalian bone marrow hematopoiesis.

**Figure 6:**
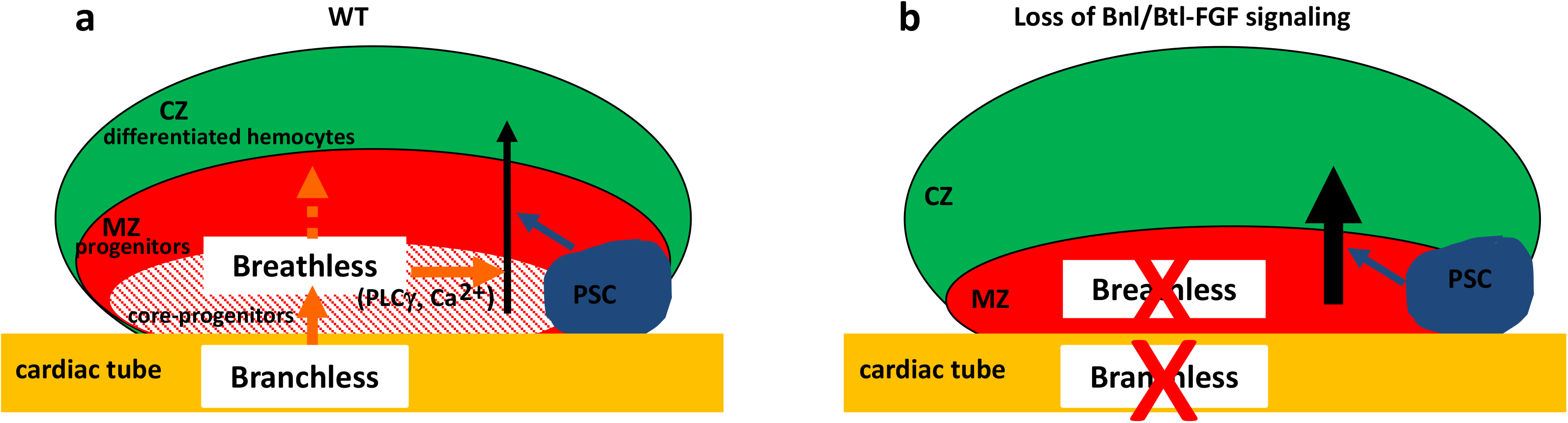
Two niches control lymph gland homeostasis. (a-b) Schematic representation of third instar larvae lymph gland anterior lobes. Progenitors and core progenitors are in red and hatched red, respectively. The cortical zone (CZ) is in green, the PSC and the cardiac tube (CT)/vascular system are in blue and orange, respectively. (a) In a wildtype (WT) lymph gland, under normal conditions the PSC, the first niche identified, regulates the maintenance of the progenitor pool except for core-progenitors (blue arrow). Here, we show that by directly acting on core-progenitors (orange arrow) the cardiac tube corresponds to a second niche present in the lymph gland. Bnl produced by cardiac cells activates its receptor Btl in progenitors. Btl-FGF activation regulates intracellular Ca^2+^ levels via PLCγ, and controls the maintenance of core-progenitors and in turn the whole progenitor pool. (b) When *bnl* or *btl* are knocked down in cardiac cells and progenitors, respectively, an increase in blood cell differentiation in the CZ is observed at the expense of the progenitor pool.

Btl-FGF signaling regulates trachea morphogenesis, which builds the insect respiratory system (Glazer and Shilo, 1991; Klambt et al., 1992; Muha and Muller, 2013; Sato and Kornberg, 2002; Sutherland et al., 1996). How the ligand Bnl diffuses from its source to activate Btl in neighboring cells remains a controversial issue. Studies performed on *Drosophila* larval Air-Sac-Primordium (ASP), using endogenous tagged versions of Bnl and Btl, brought to light a key role of long range direct cellular contacts mediated by long thin cellular extensions called cytonemes (Roy et al., 2011; Sato and Kornberg, 2002). In this process, rather than diffusing passively, Bnl produced by wing disc cells is delivered directly to ASP cells by cytonemes to activate FGF signaling (Du et al., 2018). No cytoplasmic extensions from either cardiac cells or MZ progenitors were observed so far, ruling out the delivery of Bnl from cardiac cells though long cytoplasmic extensions. Instead, both cardiac cells and MZ progenitors are embedded in a dense network of extra-cellular matrix (ECM) (Grigorian et al., 2013; Krzemien et al., 2007; Volk et al., 2014). The role of ECM components and associated cell-surface proteins, such as heparan sulfate proteoglycans (HSPGs) (Muha and Muller, 2013), in lymph gland Btl-FGF activation deserves additional investigation.

Bnl::GFP secreted by cardiac cells is detected in MZ progenitors as cytoplasmic punctate dots positive for Rab11, a marker for recycling vesicles, for Rab7, a marker for late endosomes, and for the receptor Btl. These data suggest that Bnl secreted by cardiac cells is internalized by MZ progenitors most likely through receptor-mediated endocytosis. *bnl* is transcribed in MZ progenitors and Bnl produced by these cells also contributes to lymph gland homeostasis. Taking into account the FGF dose-dependent response shown to operate in vertebrates (Ameri et al., 2010; Iyengar et al., 2007), several lymph gland sources of Bnl could be necessary to reach the threshold needed to fully activate the Btl-FGF pathway in lymph gland progenitors.

Interestingly, the Htl-FGF pathway is also required in MZ progenitors with a loss-of-function phenotype (Dragojlovic-Munther and Martinez-Agosto, 2013) opposite to that of to Btl-FGF inactivation. While Htl-FGF signaling acts through Ras and MAPK activation, we show here that Btl-FGF signaling controls intracellular calcium concentration in hematopoietic progenitors probably through PLCγ activation. By performing epistasis experiments, we further establish the absence of hierarchy between Htl-FGF and Btl-FGF signaling pathways in the MZ and that both pathways are required simultaneously to control lymph gland hematopoiesis. To our knowledge, this is the first example in which Btl and Htl are both expressed and required in the same cell population. We postulate that simultaneous regulation by the two pathways and a Bnl contribution by two separate tissue confers robustness to lymph gland hematopoiesis under normal developmental conditions and flexibility in response to environmental fluctuation. Since Htl and Btl inactivation leads to opposite lymph gland phenotypes, this raises the question of their respective downstream targets.

In vertebrates, FGF signaling controls both primitive and definitive hematopoiesis (Dzierzak and Bigas, 2018; Muha and Muller, 2013; Ornitz and Itoh, 2015). Additional studies indicate that FGFR1 in adult HSPCs is activated during hematopoietic recovery following injury, in order to stimulate HSPC proliferation and mobilization (Zhao et al., 2012). Furthermore, FGF2 facilitates HSPC expansion by amplifying mesenchymal stem cells, a niche cell type (Itkin et al., 2013; Itkin et al., 2012). Thus, in vertebrate adult bone marrow, the FGF pathway plays a major role in the control of hematopoiesis both under steady state conditions and in response to an immune stress. However, deciphering how FGF controls hematopoiesis in bone marrow remains an arduous task since many FGF ligands and receptors are expressed in HSPC and/or niche cells and redundancy and compensation mechanisms between different FGF members can occur (Haas et al., 2018). Given the high conservation of signaling pathways between *Drosophila* and mammals, the low genetic redundancy in *Drosophila* and the striking similarities between mammalian bone marrow and fly lymph gland, there is promise that our newly identified regulation of FGF signaling in the lymph gland will shed light on the complex regulation of FGF signaling in mammalian bone marrow.

## Materials and methods

### Fly strains

*w^1118^* (wild type, *WT*), *UAS-mCD8-GFP* and PG125*dome-gal4* (Krzemien et al., 2007), *antp-gal4* (Mandal et al., 2007)*, handΔ-gal4* (Morin-Poulard et al., 2016) and *NP1029-gal4* (Monier et al., 2005)*. handΔ-gal4* corresponds to the 3^rd^ intron of *hand* deleted from the specific visceral mesoderm enhancer ((Popichenko et al., 2007) and Laurent Perrin personal communication). Lymph gland mcd8-GFP expression patterns under *handΔ-gal4* and *NP1029-gal4 drivers* in L1, L2 and L3 larvae are given in Figure 1- figure supplement 1. The *handΔ-gal4* is expressed in all cardiac cells, whereas the NP1029-*gal4* is expressed in all cardiac cells except those that are expressing *seven-up* (Monier et al., 2005). Strains used are BcGFP (Tokusumi et al., 2009), *bnl^P2^* (Sutherland et al., 1996), *hhF4-GFP* (Tokusumi et al., 2010), *domeMESO-LacZ* (Krzemien et al., 2007), *domeMESO-Gal4* (Louradour et al., 2017), *UAS-Bnl::GFP* (Lin and Affolter, 2009), *UAS-btl^CA^* on II or III (Pares and Ricardo, 2016), *UAS-Bnl* (Jarecki et al., 1999). *Ubiquitin-rab11cherryFP* (Y. Bellaiche), *bnl:GFP^endo^* and *btl:cherry^endo^* knock-in alleles (Du et al., 2018). Other strains were provided by the Bloomington (BL) and the Vienna (VDRC) *Drosophila* RNAi stock centers: *btl^dev1^* (BL4912), *Sar1-RNAi* (BL 32364, (Cook et al., 2017)), *UAS-CaMKII* (BL 29662), *UAS-GCaMP3* (BL32116), *UAS-IP3R* (BL30741), *sl-RNAi* (BL32385 and BL35604), *sl^2^* (BL724). The list of RNAi lines used for the functional screen is given in Figure 1-figure supplement 2. For RNAi treatments, *UAS-Dicer 2* was introduced and at least two RNAi lines per gene were tested. Controls correspond to Gal4 drivers crossed with *w^1118^*. In all experiments, crosses and subsequent raising of larvae until late L1/early L2 stage were performed at 22°C, before shifting larvae to 29°C until their dissection at the L3 stage. For gal*80^ts^* experiments, crosses were initially maintained at 18°C (permissive temperature) for 3 days after egg laying, and then shifted to 29°C until dissection.

### RNAi screen

Antenapedia (Antp) immunostaining was revealed with the ABC kit from Abcam. The images were collected with a Nikon epifluorescence microscope. PSC cell numbers were counted manually using Fiji multi-point tool software. The BcGFP and anterior lobe areas were measured. Crystal cell index corresponds to BcGFP area/anterior lobe area. 2 RNAi lines per gene were tested when available, and at least 15 lymph glands were analyzed per genotype.

### Generation of DomeMESO-RFP transgenic lines

The domeMESO sequence from *pCasHs43domeMESO-lacZ* (Rivas et al., 2008) was sub-cloned into pENTR Directional, following the experimental procedure of the TOPO®Cloning Kit from Invitrogen. The resulting plasmid was used to generate *domeMESO-RFP* transgenic flies using attP/attB technology (Bischof et al., 2007). The *Drosophila* line was created by integration at *attP*-68A4 (III) sites.

### Antibodies and immunostaining

Lymph glands were dissected and processed as previously described (Krzemien et al., 2007). Antibodies used were mouse anti-Col (1/100) (Krzemien et al., 2007), chicken anti-βgal (1/1000, Abcam), rabbit anti-RFP (1/40 000, Rockland Immunochemicals), chicken anti-GFP(1/500, Abcam), mouse anti-Antp (1/100, Hybridoma Bank), mouse anti-Hnt (1/100, Hybridoma Bank); mouse anti-P1 (1/30, I. Ando, Institute of Genetics, Biological Research Center of the Hungarian Academy of Science, Szeged, Hungary), mouse anti-proPO (1/100, T.Trenczel, Justus-Liebig-University Giessen, Giessen, Germany). Secondary antibodies were Alexa Fluor-488 and −555 conjugated antibodies (1:1000, Molecular Probes) and goat anti-Chicken Alexa Fluor-488 (1/800; Molecular Probes). Nuclei were labeled with TOPRO3 (Thermo Fisher Scientific). Immunostainings were performed as previously described (Louradour et al., 2017). For detecting bnl:GFP^endo^ and btl:cherry^endo^ immunostainings were performed with anti-GFP and anti-RFP, respectively.

### *In situ* hybridization

The protocol was as described in (Oyallon et al., 2016). For fluorescent *in situ* hybridization we used digoxigenin-labelled *tep4* and *bnl* probes. For revelation, samples were incubated with sheep-anti-DIG (1/1000, Roche) followed by biotinylated donkey-anti-sheep (1/500, Roche). ABC kit from Vector Laboratory was used followed by fluorescent tyramide staining (Alexa fluor 555 or 488 conjugated tyramide from Molecular Probes). The *bnl* probe was transcribed in *vitro* using T7 RNA polymerase II, from PCR-amplified DNA sequences. Pairs of primers were used and the sequence in italics corresponds to the T7 RNA-Pol II fixation site. For *bnl*: primer 1: GCCATGGACAACAACTTGAC/*ATGAATTCTAATACGACTCACT ATAGGG*CGTCGTTACGGTCCAGATTG; primer 2: GCAAGGCCAACAAGAAGAAG/ *ATGAATTCTAATACGACTCACTATAGGG*CCTGGTCGTTATCCTGATCC.

### Quantification of PSC cell numbers

In all experiments, genotypes were analyzed in parallel and quantified. PSC cells were counted manually using Fiji multi-point tool software. Statistical analyses (Mann–Whitney nonparametric test) were performed using GraphPad Prism 5 software.

### Blood cell and progenitor quantification

Crystal cells were visualized by either BcGFP or immunostainings with antibodies against proPO or Hnt. Plasmatocytes were labelled by P1 immunostainings. *DomeMESO-RFP* and *DomeMESO-GFP* were expressed in MZ progenitors, whereas MZ core progenitors were labelled by either *tep4 in situ* hybridization or Col immunostainings. Optimized confocal sections were performed on Leica SPE or SP8 microscopes for 3D reconstruction. The numbers of crystal cells, plasmatocytes and progenitors stained and anterior lobe volume (in μm^3^) were measured using Volocity 3D Image Anaysis software (PerkinElmer). Crystal cell index: (crystal cell number/anterior lobe volume)x100000; plasmatocyte and progenitor index: (plasmatocyte or progenitor volume/anterior lobe volume)x100. At least 15 anterior lobes were scored per genotype, and experiments were reproduced at least three times. Statistical analyses (Mann–Whitney nonparametric test) were performed using GraphPad Prism 5 software. Since the number of lymph gland differentiated blood cells fluctuates depending on the larval stage, and to limit discrepancies in all the experiments, genotypes were analyzed in parallel.

### Quantification of hhF4-GFP and UAS-GCaMP3 intensity

Optimized confocal sections were performed on Leica SPE or SP8 microscopes for 3D reconstruction. For hhF4-GFP, the sum intensities for GFP per PSC labelled by Col and each PSC volume (in μm3) were measured using Volocity 3D Image Anaysis software (PerkinElmer). The intensity of hhF4-GFP corresponds to the sum intensity of hhF4-GFP/the PSC volume. For GCaMP3, the sum intensities for GFP per lymph gland primary lobe labelled by TOPRO and each primary lobe volume (in μm3) were measured using Volocity 3D Image Anaysis software. The intensity of GCaMP3 corresponds to the sum intensity of GCaMP3/per lymph gland primary lobe volume. At least 15 anterior lobes were scored per genotype, and experiments were reproduced at least three times. Statistical analyses (Mann–Whitney nonparametric test) were performed using GraphPad Prism 5 software.

### Quantification of the diffusion in the MZ of cytoplasmic Bnl::GFP dots, in hand>Bnl::GFP and hand>Bnl::GFP>sar1-RNAi genetic contexts

Optimized lymph gland confocal sections were obtained with a Leica SP8 microscope for 3D reconstruction. The maximum projection of 10 slices chosen in the middle of the stack was performed. A parallelepiped with a larger corresponding to 4 nuclei diameter, a width of 10 confocal slices a length corresponding to the distance from the CT to the CZ was designed. Along the length, the parallelepiped was subdivided into 11 sub parallelepipeds of similar size. The number of Bnl::GFP granules per sub parallelepiped (called interval in Figure 4g legend) was counted. Spot detector plugin from ICY software (http://icy.bioimageanalysis.org/) was used to quantify the number of Bnl::GFP dots per sub parallelepiped.

### Sample size

n corresponds to the number of anterior lobes analyzed. **Figure 1:** In e, for *handΔ>* n=61 and n=26 for *handΔ>ilp6-RNAi*. In h, for *handΔ>* n=61 and n=24 for *handD>ds-RNAi*. In k, *handΔ>* n=61 and n=24 for *handΔ>pvf3-RNAi*. In f, for *handΔ>* n=19 and n=27 for *handΔ>ilp6-RNAi*. In i, for *handΔ>* n= 26 and n=21 *for handΔ>ds-RNAi*. In l, *handΔ>* n=26 and n=26 for *handΔ>pvf3-RNAi*. In o, for *handΔ>* n=14 and n=10 for *handΔ>pvf3-RNAi*. **Figure 2:** in d, n=12 for WT and n=10 for *bnl^P2^/+.* In i, *handΔ>* n=22, *handΔ>bnl-RNAi* n=13, *HandΔ>bnl* n=38 and *handΔ>bnl;bnl-RNAi* n=10. In l, for *handΔ>* n=24 and n=15 for handΔ>bnl-RNAi. In o, for handΔ> n=25 and n=28 in *handΔ>bnl-RNAi*. For r, *handΔ>* n=36 and n=22 for *handΔ>bnl-RNAi*. In u, for handΔ> n=16 and n=18 in *handΔ>bnl-RNAi*.

**Figure 3:** In d, n=45 for *WT* and n=30 for *btl ^dev1^/+.* In g, dome> n=93, *dome>btl-RNAi* n=23, *dome>btl^CA^* n=12 and *dome>btl^CA^;btl-RNAi* n=20. In j, n=27 for *dome>* and n=21 for *dome>btl-RNAi*. In m, n=20 for *dome>* and n=24 for *dome>btl-RNAi*. In p, n=16 for *dome>*, n=13 for *dome>htl^DN^*. n=18 for *dome>btl-RNAi* and n=29 for *dome> dome>htl^DN^>btl-RNAi*.

**Figure 4:** In k: *handΔ>* n=27, *handΔ>sar1-RNAi* n=14 and *handΔ>sar1-RNAi*; *Bnl//GFP* n=17.

**Figure 5:** In c, n =41 for *dome>* and n=29 for *dome>btl-RNAi*. In h, for *dome>* n=35, n= 33 for *dome>btl-RNAi*, n=35 for *dome>CaMKII* and n=26 for *dome>CaMKII*; *btl-RNAi*. In l, *dome>* n=27 and n=30 in *dome>sl-RNAi*. In m, *WT* n=20 and n=17 in *sl^2^*. In n, *sl^2^, dome> n*=20 and n=21 in *sl^2^; dome>btlCA*.

### Replicates

**Figure 2:** d, i) 3 independent experiments were performed and quantified. One is shown. l) 2 independent experiments were performed and quantified. One is shown. o) 3 independent experiments were performed and quantified. One is shown. r) 2 independent experiments were performed and quantified. One is shown. u) 2 independent experiments were performed and quantified. One is shown. **Figure 3:** d, g) 3 independent experiments were performed and quantified. One is shown. j) 2 independent experiments were performed and quantified. One is shown. m) 3 independent experiments were performed and quantified. One is shown. p) 2 independent experiments were performed and quantified. One is shown **Figure 4:** k) 2 independent experiments were performed and quantified. One is shown. **Figure 5 c, h, l-n**) 2 independent experiments were performed and quantified. One is shown.

### Drosophila genetics

Fly crosses for each figure:

Figure 1c-c’, m: *handΔ,UAS-dcr2; BcGFP crossed with w^1118^*; Figure 1d, d’ *: handΔ,UAS-dcr2; BcGFP crossed with UAS-ilp6-RNAi*; Figure 1g, g’ *: handΔ,UAS-dcr2; BcGFP crossed with UAS-ds-RNAi*; Figure 1j, j’, n *: handΔ,UAS-dcr2; BcGFP crossed with UAS-pvf3-RNAi.* Figure 2a-a”: *domeMESO-GFP crossed with w^1118^*; Figure 2b: *w^1118^*; Figure 2c: *bnl^P2^/TM6B crossed with w^1118^*; Figure 2e: *handΔ,UAS-dcr2; BcGFP crossed with w^1118^;* 2f: *handΔ,UAS-dcr2; BcGFP crossed with UAS-bnl-RNAi;* 2g: *handΔ,UAS-dcr2; BcGFP crossed with UAS-bnl;* 2h: *handΔ,UAS-dcr2; BcGFP crossed with UAS-bnl;UAS-bnl-RNAi; 2j: handΔ,UAS-dcr2; domeMESO-RFP crossed with w^1118^; 2k: handΔ,UAS-dcr2; domeMESO-RFP crossed with UAS-bnl-RNAi; 2m and p: handΔ,UAS-dcr2 crossed with w^1118^; 2n and q: handΔ,UAS-dcr2 crossed with UAS-bnl-RNAi. s: handΔ,UAS-dcr2; bnl:GFP^endo^ crossed with w^1118^; t: handΔ,UAS-dcr2; bnl:GFP^endo^ crossed with UAS-bnl-RNAi.*

Figure 3a-a”: *domeMESO-GFP* crossed *with btl:mcherry^endo^*; 3b and k: *PG125dome-gal4,UAS-dcr2* crossed with *w^1118^*; c *btl^dev1^/TM6B* crossed with *w^1118^*; e and i: *PG125dome-gal4,UAS-dcr2* crossed with *UAS-btl-RNAi; f: PG125dome-gal4,UAS-dcr2* crossed with *UAS-btl^CA^; UAS-btl-RNAi; h: PG125dome-gal4,UAS-dcr2*; *DomeMESO-LacZ* crossed with *w^1118^; i: PG125dome-gal4,UAS-dcr2*; *DomeMESO-LacZ* crossed with *UAS-btl-RNAi*; *n: PG125dome-gal4,UAS-dcr2* crossed with *UAS-htl^DN^*; *o: PG125dome-gal4,UAS-dcr2* crossed with *UAS-htl^DN^; UAS-btl-RNAi.*

Figure 4a: *handΔ,UAS-dcr2 crossed with UAS-bnl::GFP; 4b: handΔ,UAS-dcr2; btl:mcherry^endo^ crossed with UAS-Bnl::GFP; 4c: handΔ,UAS-dcr2; UAS-bnl::GFP crossed with ubi-rab11::mcherry;* 4d: *handΔ,UAS-dcr2 crossed with UAS-bnl::GFP*; 4e: *handΔ,UAS-dcr2;UAS-Bnl::GFP crossed* with *w^1118^;* 4f: *handΔ,UAS-dcr2; Bnl::GFP crossed with UAS-sar1-RNAi*; *4h: handΔ,UAS-dcr2; BcGFP crossed with w^1118^; 4i: handΔ,UAS-dcr2; BcGFP crossed with:UAS-sar1-RNAi; 4j: handΔ,UAS-dcr2; BcGFP crossed with UAS-sar1-RNAi; UAS-bnl::GFP.*

Figure 5a: *PG125dome-gal4,UAS-dcr2 crossed with UAS-GCaMP3*; 5b: *PG125dome-gal4,UAS-dcr2 crossed with UAS-GCaMP3; UAS-btl-RNAi*; 5d: *PG125dome-gal4,UAS-dcr2 crossed with w^1118^*; 5e:*PG125dome-gal4,UAS-dcr2 crossed with UAS-btl-RNAi*; 5f:*PG125dome-gal4,UAS-dcr2 crossed with UAS-CaMKII*; 5g:*PG125dome-gal4,UAS-dcr2 crossed with UAS-CaMKII; UAS-btl-RNAi*; 5i:*PG125dome-gal4,UAS-dcr2 crossed with UAS-sl-RNAi*; 5j: *sl^2^; 5k: sl^2^; dome-gal4 crossed with sl^2^; UAS-btl-*CA.

Figure 1-figure supplement 1a-c: *handΔ,UAS-dcr2 crossed with UAS-mcd8GFP; d-f:NP1029,UAS-dcr2 crossed with UAS-mcd8GFP.*

Figure 2 - figure supplement 1a*: BcGFP crossed with w^1118^*; b: *hml-gal4*, *UAS-mcd8GFP* crossed *with w^1118^*; c: *pcol-gal4, UAS-mcd8-GFP crossed with w^1118^*; *d: handΔ,UAS-dcr2 crossed with UAS-mcd8-GFP; e: handΔ,UAS-dcr2* crossed *UAS-mcd8-GFP; UAS-bnl-RNAi;* f: *handΔ,UAS-dcr2* crossed *with w^1118^; g: handΔ,UAS-dcr2* crossed *with UAS-bnl-RNAi; i: handΔ,UAS-dcr2 crossed with w^1118^; j: handΔ,UAS-dcr2 crossed with UAS-bnl-RNAi 34572; l: NP1029, UAS-dcr2* crossed *with w^1118^; m: NP1029, UAS-dcr2* crossed *with UAS-bnl-RNAi; o: handΔ,UAS-dcr2;tub-gal80^ts^* crossed *with w^1118^; p: handΔ,UAS-dcr2;tub-gal80^ts^* crossed *with UAS-bnl-RNAi.*

Figure 2-figure supplement 2a: *PG125dome-gal4,UAS-dcr2* crossed with *w^1118^*; b: *PG125dome-gal4,UAS-dcr2* crossed with *UAS-bnl-RNAi; d: PG125dome-gal4,UAS-dcr2* crossed with *w^1118^; e: PG125dome-gal4,UAS-dcr2* crossed with *UAS-bnl-RNAi; g: handΔ,UAS-dcr2* crossed with *w^1118^; h: handΔ,UAS-dcr2* crossed with *UAS-bnl; UAS-bnl-RNAi; j: handΔ,UAS-dcr2* crossed with *w^1118^; k: handΔ,UAS-dcr2* crossed with *UAS-bnl-RNAi; m: hhF4-GFP; handΔ,UAS-dcr2 crossed with w^1118^; n: hhF4-GFP; handΔ,UAS-dcr2* crossed with *UAS-bnl-RNAi.*

Figure 3-figure supplement 1; a: *BcGFP* crossed with *btl:cherry^endo^*; b: *hml-gal4, UAS-mcd8-GFP* crossed with *btl:cherry^endo^*; c: *pcol-gal4, UAS-mcd8-GFP* crossed with *btl:cherry^endo^*; d: *handΔ,UAS-dcr2; BcGFP* crossed with *w^1118^*; e*: handΔ,UAS-dcr2; BcGFP crossed with UAS-btl-RNAi*; g*: handΔ,UAS-dcr2 crossed with w^1118^; h: handΔ,UAS-dcr2 crossed with UAS-btl-RNAi; j and m: PG125dome-gal4,UAS-dcr2* crossed with *w^1118^*; k and n: *PG125dome-gal4,UAS-dcr2* crossed with *UAS-btl-RNAi.*

Figure 4-figure supplement1a: *handΔ, UAS-dcr2* crossed with *w^1118^*; *b: handΔ, UAS-dcr2* crossed with *UAS-sar1-RNAi; d and g: NP1029, UAS-dcr2* crossed with *w^1118^*; *e and h: handΔ, UAS-dcr2* crossed with *UAS-sar1-RNAi.*

Figure 5-figure supplement 1: a: *PG125dome-gal4,UAS-dcr2* crossed with *UAS-GCaMP3; b: PG125dome-gal4,UAS-dcr2* crossed with *UAS-GCaMP3;UAS-btl-RNAi; d: tep4-gal4,UAS-dcr2* crossed with *w^1118^; e: tep4-gal4,UAS-dcr2* crossed with *UAS-btl-RNAi; f: tep4-gal4,UAS-dcr2* crossed with *UAS-IP3R; g: tep4-gal4,UAS-dcr2* crossed with *UAS-IP3R; UAS-btl-RNAi.*

## Acknowledgements

We thank M. Affolter, Y. Bellaiche, J. Casanova, J. Colombani, L. Du, M. Freeman, M. Grammont, M. A. Krasnow, A. Paululat, L. Perrin, S. Ricardo, S. Roy, T. Tanaka, P. Thérond, Bloomington and Vienna Stock Center and the TRiP at Harvard Medical School for fly strains; I. Ando, A. Moore and T. Trenczek for antibodies; L. Bataillé, A. Davy, G. Lebreton, M. Meister, C. Monod, B. Monnier and A. Vincent, for critical reading of the manuscript. We are grateful to B. Ronsin and S. Bosch for assistance with confocal microscopy (Plateforme TRI); J. Favier, V. Nicolas and A. Destenable for fly culture. Research in the authors’ laboratory is supported by the CNRS, University Toulouse III, Ministère de la Recherche (ANR « programme blanc »), ARC (Association pour la Recherche sur le Cancer), La Ligue contre le Cancer 31, La Société Française d’Hématologie (SFH) and FRM (Fondation pour la Recherche Médicale).

## Author contributions

Conceived and designed the experiments: M.D.L. and M.C. Performed the experiments: M.D.L., I.M.P., Y.T. and N.V. Analyzed the data: M.D.L., I.M.P., N.V. and M.C. Contributed reagents/materials/analysis tools: I.M.P. Wrote the paper: M.D.L and M.C.

---

**Figure1-figure supplement 1.**
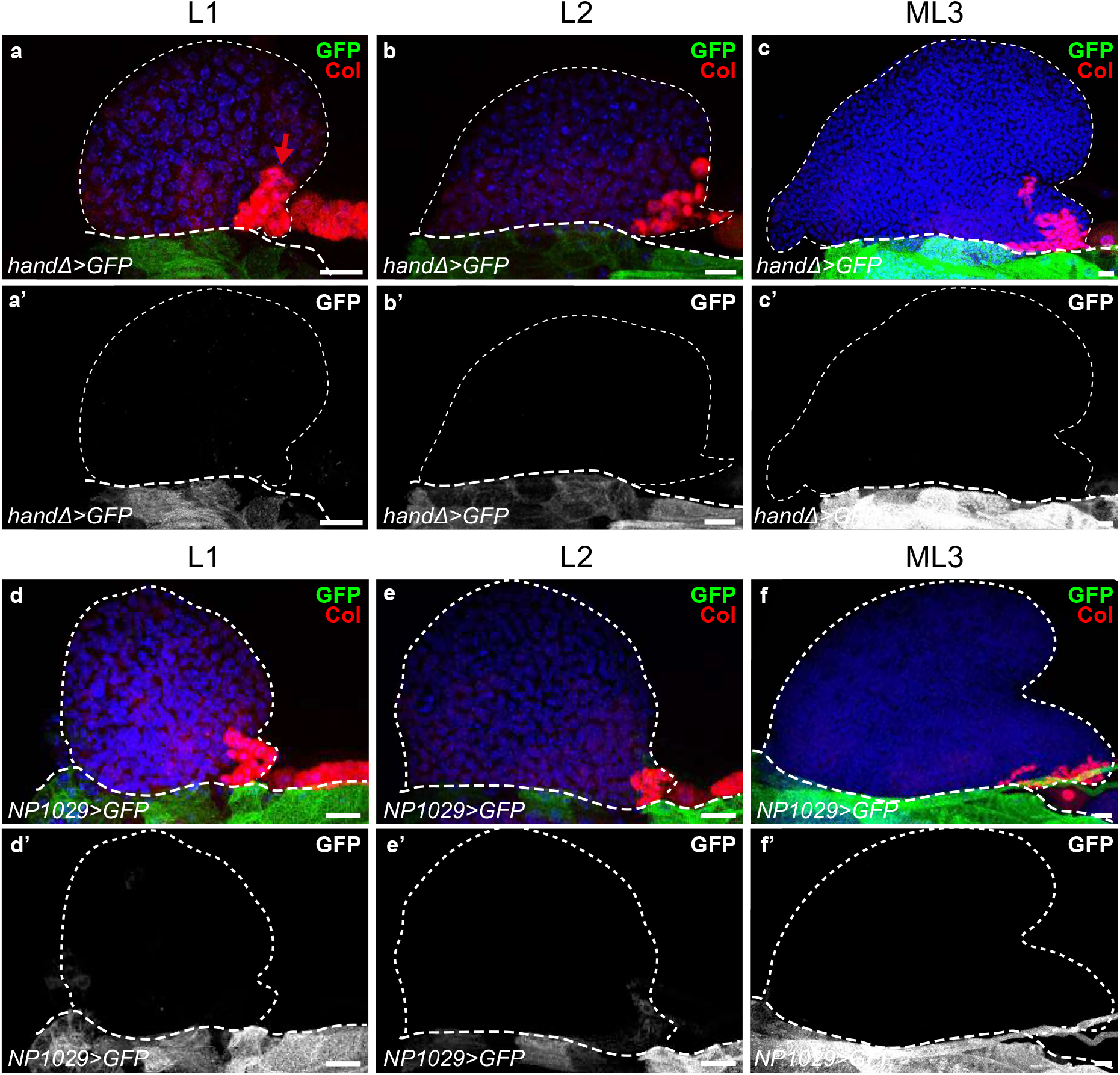
Expression pattern of *handΔ-gal4* and *NP1029-gal4* driver in lymph glands during larval development. (a-c’) *handΔ-gal4>mcd8-GFP* is green in (a-c) and white in (a’-c’). (d-f’) *NP1029-gal4>mcd8-GFP* is green in (d-f) and white in (d’-f’). The PSC is labelled by Col (red and arrow in a); thin and thick dashed lines indicate the contours of the lymph gland and the cardiac tube, respectively.

**Figure1-figure supplement 2.**
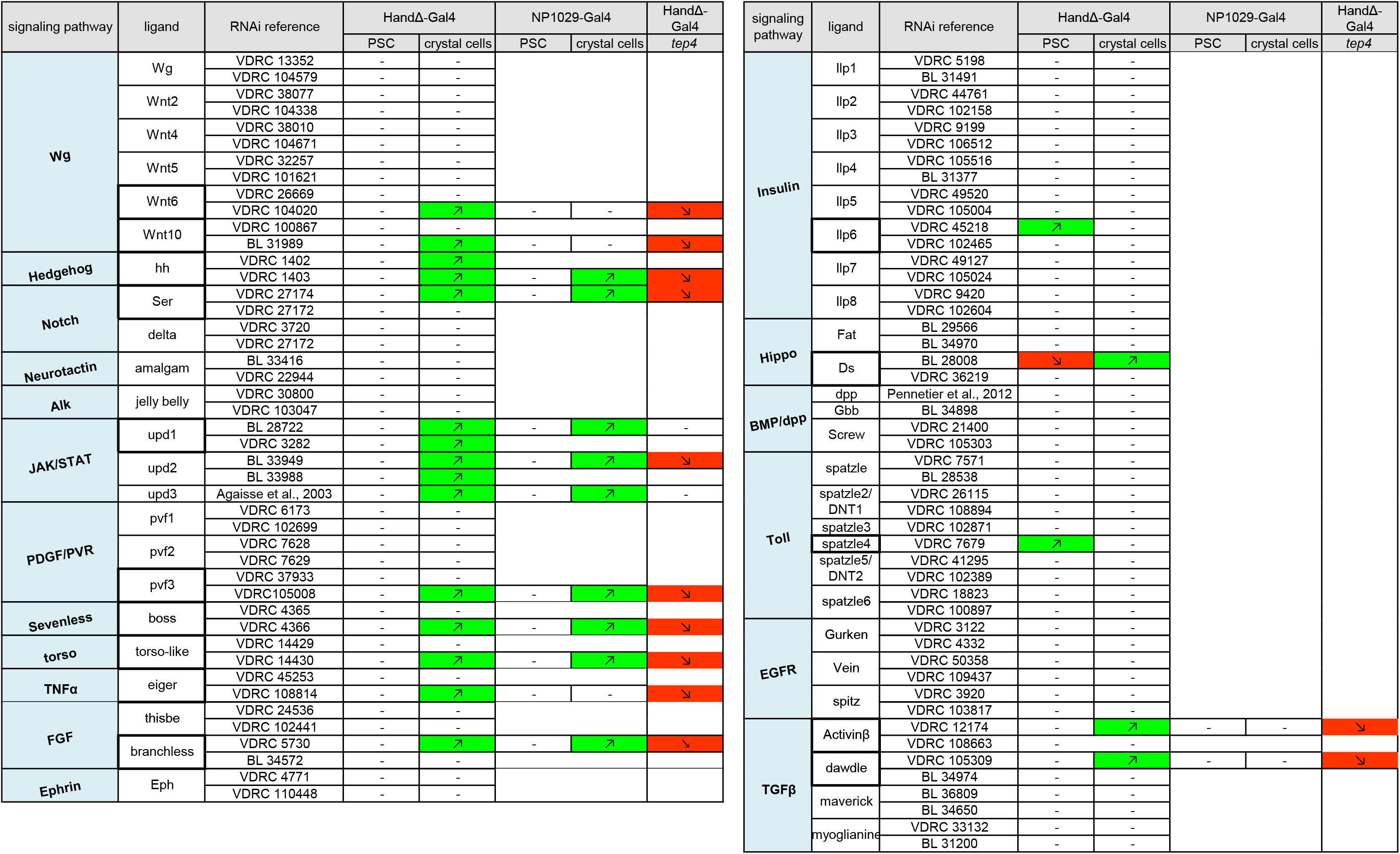
Results of the RNAi ligand screen. RNAi was expressed in cardiac cells by using the *handΔ-gal4* and/or *NP1029-gal4* driver. Crystal cells were labeled by BcGFP, PSC cells were immune-stained with Antp antibody, and to visualize the core progenitors *tep4 in situ* hybridization was performed. In most cases, 2 RNAi lines were tested per ligand, and at least 15 lymph glands per RNAi were analyzed. Crystal cell index and PSC cell number were established. The green and red colored boxes indicate an increase and a decrease, respectively, compared to the control. Black dashes indicate that no difference was observed compared to the control. A white box indicates that this condition was not tested. Most RNAi lines that gave a modification in crystal cell index with the *handΔ-gal4* driver were also analyzed with another cardiac cell driver *NP1029-gal4,* and proPO antibody immunostainings were performed to visualize crystal cells. Finally, for all RNAi lines that led to a defect in crystal cell differentiation with the *handΔ-gal4* driver, *tep4 in situ* hybridizations were performed and the *tep4* index was established.

**Figure2-figure supplement 1.**
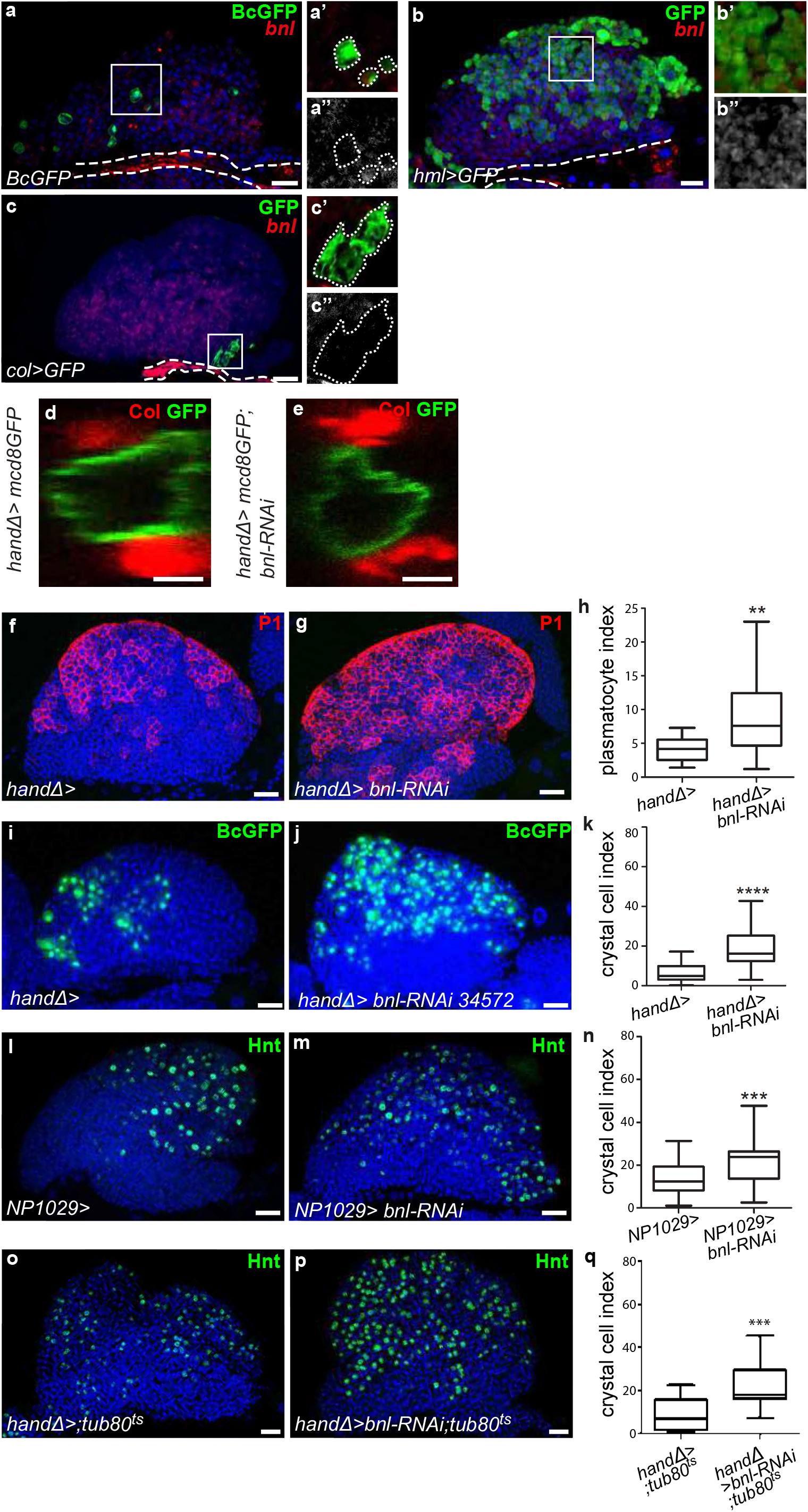
The ligand Bnl in cardiac cells controls lymph gland hemocyte differentiation homeostasis. (a-c”) A maximum projection of 5 confocal lymph gland sections, *bnl* is in red. (a,b,c) and white in (a”,b”,c”). BcGFP (green) labels crystal cells. (a’-a”) An enlarged view, crystal cells are green (a’) and *bnl* is white (a”). (b-b”) *hml>GFP* (green) labels differentiating hemocytes in the CZ. (b’-b”) An enlarged view, differentiating hemocytes are green (b’) and *bnl* is white (b”). (c-c”) *col>GFP* (green) labels PSC cells. (c’-c”) An enlarged view, the PSC is green (c’) and *bnl* is white (c”). White large dashed line indicates the cardiac tube contour. (d, e) Cardiac cells express mcd8-GFP (*handΔ>mcd8GFP*, green). A transversal section is shown. PSC cells are labelled by Col (red). No difference in cardiac tube morphology is observed when *bnl* is knocked down in cardiac cells at the larval stage (e) compared to the control (d). (f, g) Plasmatocytes (red) are labelled by P1 antibody. An increase in plasmatocyte number is observed when *bnl* expression is decreased in cardiac cells (g) compared to the control (f). (h) Plasmatocyte index. (i-j) Crystal cells (GFP, green) express the Black-cell GFP (BcGFP) marker. An increase in crystal cell number (j) is observed when *bnl* expression is decreased in cardiac cells compared to the control (i), when another *bnl* RNAi is used (j) or when another cardiac cell driver NP1029 is used (l, m). (k, n) Crystal cell index. (o, p) Hnt labels crystal cells. Compared to the control (o) crystal cell number is increased when *bnl* is knocked down in cardiac cells from the L2 stage by using the conditional Gal4/Gal80^ts^ system (p). (q) Crystal cell index.

**Figure2-figure supplement 2.**
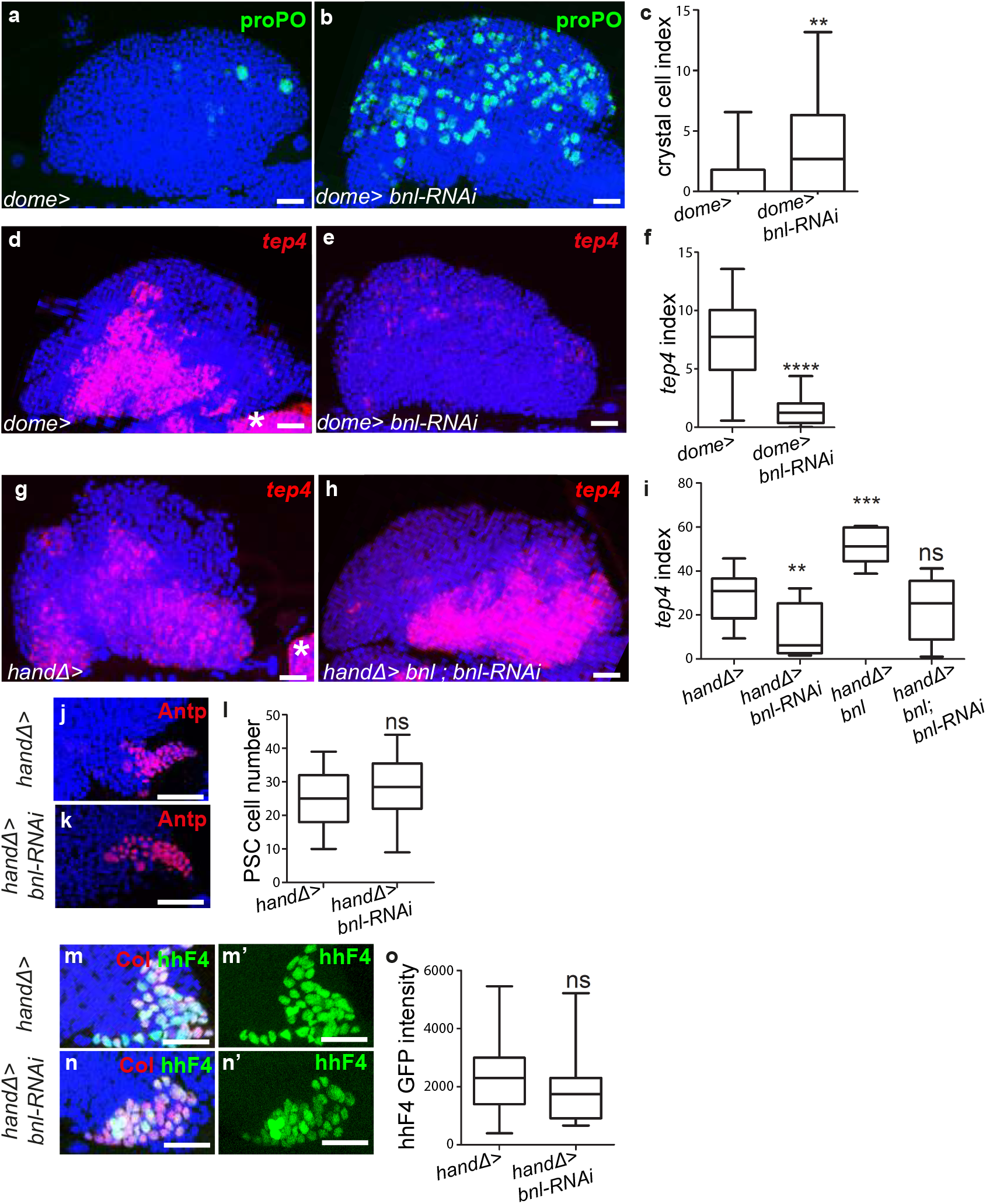
Whereas the ligand Bnl in cardiac cells does not control PSC cells, it is required in MZ progenitors to regulate lymph gland homeostasis. (a, b) Crystal cells are labeled by proPO antibody (green). An increase in crystal cell number is observed when *bnl* expression is decreased in MZ progenitors (b) compared to the control (a). (c) Crystal cell index. (d-e, g-h) *tep4* (red) is expressed in MZ progenitors. Compared to the control (d) barely detectable levels of *tep4* are observed when *bnl* is knocked down in MZ progenitors (e). Co-expression of *bnl* and *bnl* RNAi in cardiac cells restores WT levels of *tep4* expression (h and i) compared to the control (g). (f, i) *tep4* index. (j, k) PSC cells are labelled by Antp (red) antibody. No difference in PSC cell numbers is observed when *bnl* is knocked down in cardiac cells (k) compared to the control (j). (l) Quantification of PSC cell number. (m-n’) hhF4-GFP labels PSC cells (GFP, green). No difference in hhF4-GFP expression is observed when *bnl* is knocked down in cardiac cells (n) compared to the control (m). (o) Quantification of hhF4-GFP intensity.

**Figure3-figure supplement 1.**
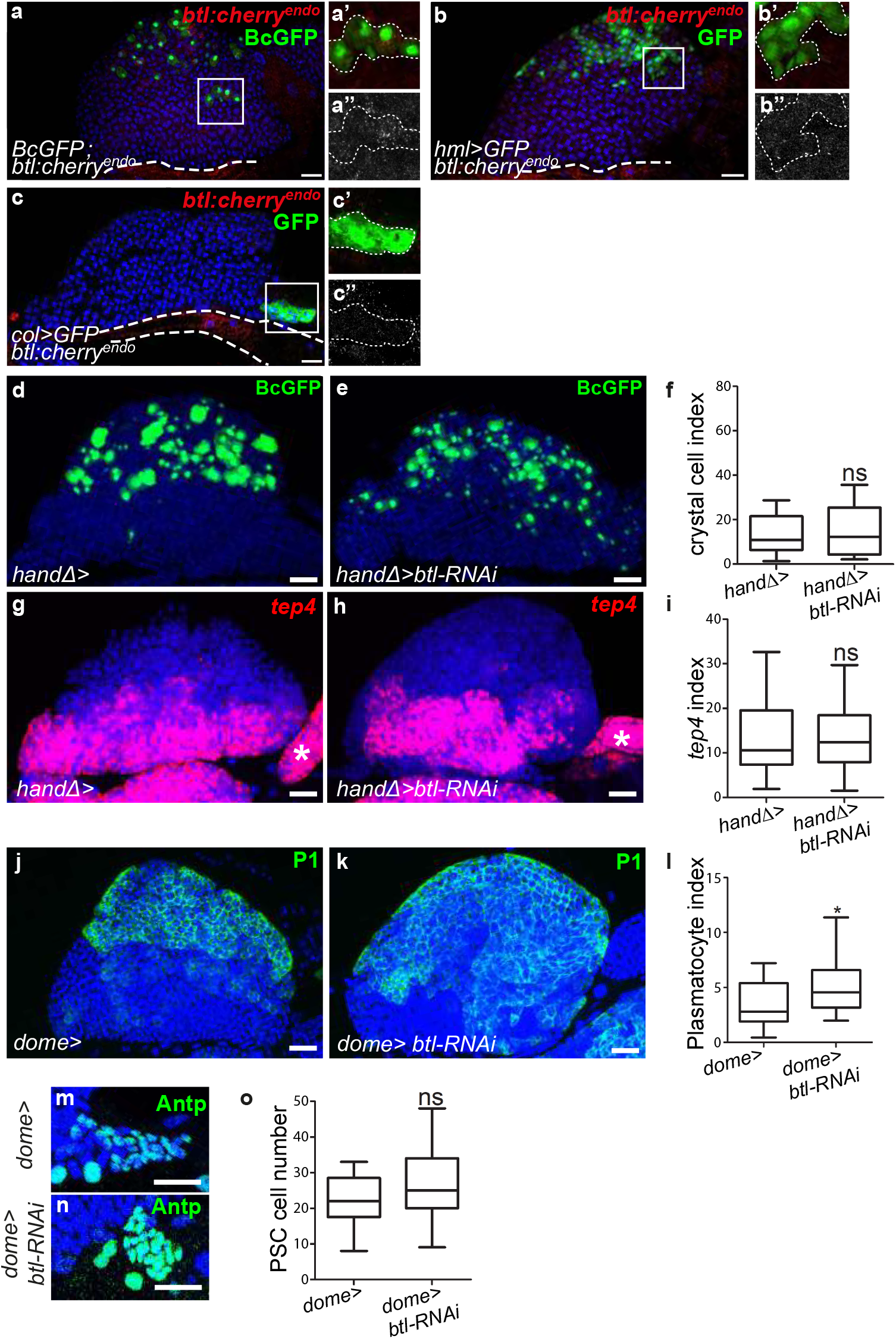
The Btl receptor in progenitors controls lymph gland hemocyte differentiation without affecting PSC size. (a-c”) A maximum projection of 5 confocal lymph gland sections of larvae expressing *btl:cherry^endo^* (red). (a-a’) BcGFP (green) labels crystal cells. (a’-a”) An enlarged view, crystal cells are green (a’) and *btl:cherry^endo^* is white (a”). (b-b”) *hml>GFP* (green) labels differentiating hemocytes in the CZ. (b’-b”) An enlarged view, differentiating hemocytes are green (b’) and *btl:cherry^endo^* is white (b”). (c-c”) *Col>GFP* (green) labels PSC cells. (c’-c”) An enlarged view, the PSC is green (c’) and *btl:cherry^endo^* is white (c”). (a, b, c) White large dashed line indicates the cardiac tube contour. (d, e) BcGFP (green) labelled crystal cells. No significant difference in crystal cell number is observed when *btl* expression is decreased in cardiac cells (e) compared to the control (d). (f) Crystal cell index. (g, h) Decreasing *btl* expression in cardiac cells (h) does not change *tep4* expression (red) compared to the control (g). (i) *tep4* index. (j, k) An increase in plasmatocyte number (labeled by P1 antibodies, green) is observed when *btl* expression is decreased in progenitors (k) compared to the control (j). (l) Plasmatocyte index. (m, n) PSC cells are labelled by Antp (green) antibody. No difference in PSC cell number is observed when *btl* is knocked down in progenitors (n) compared to control (m). (o) Quantification of PSC cell number.

**Figure4-figure supplement 1.**
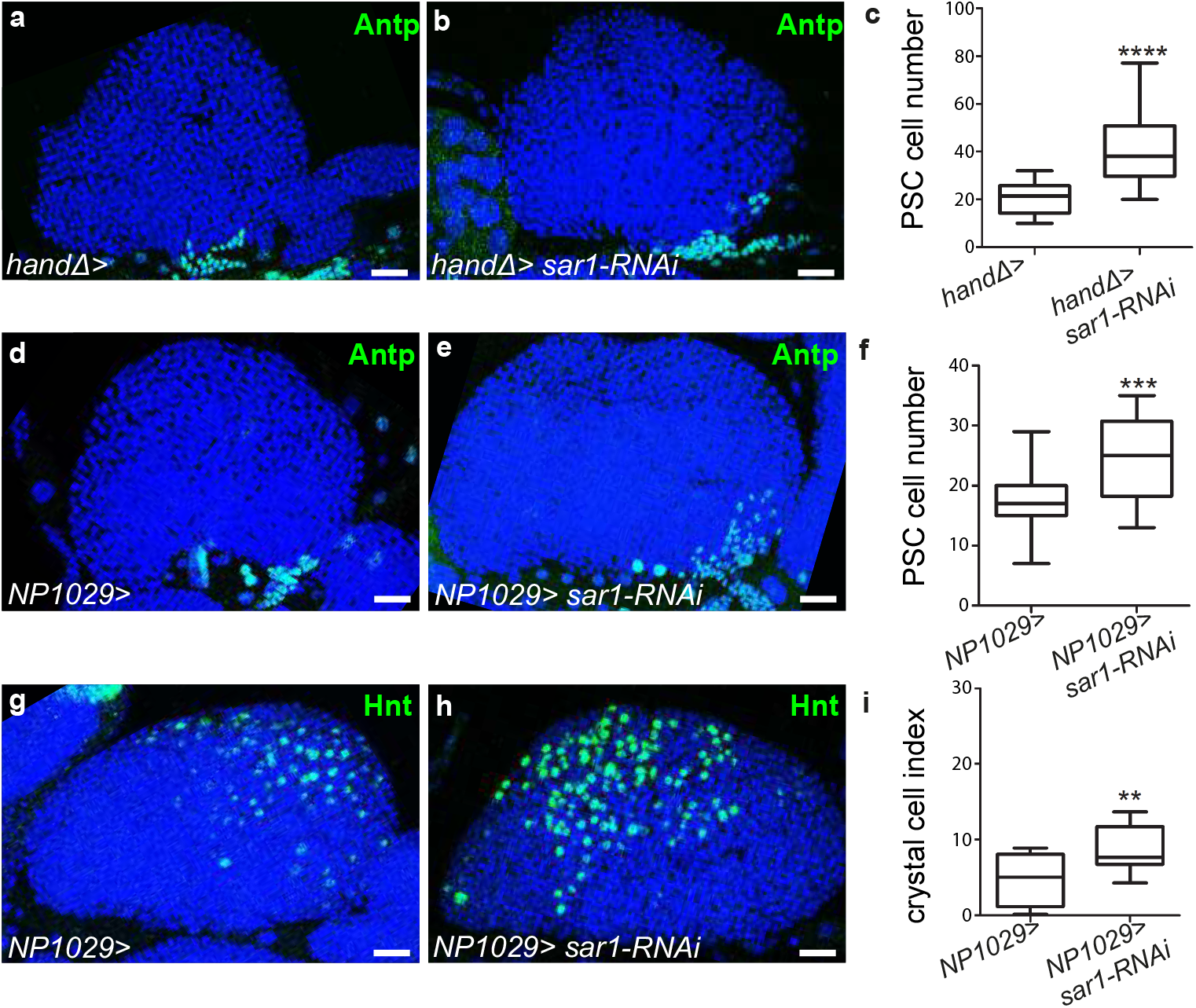
Knocking down *sar1* in cardiac cells impairs crystal cell differentiation and increases PSC size. (a-b, d-e) PSC cells are labelled with Antp (green) antibody. An increase in PSC cell numbers is observed when *sar1* is knocked down in cardiac cells using the *handΔ* driver (b) compared to the control (a). (c) Quantification of PSC cell numbers. (d-f) An increase in PSC cell number is observed when *sar1* is decreased in cardiac cells using the *NP1029* driver (e) compared to the control (d). (f) PSC cell numbers. (g, h) Crystal cells are labelled with Hnt (green) antibody. An increase in crystal cell number is observed when *sar1* is knocked down in cardiac cells using *NP1029*, another cardiac cell driver (h) compared to the control (g). (i) Crystal cell index.

**Figure5-figure supplement 1.**
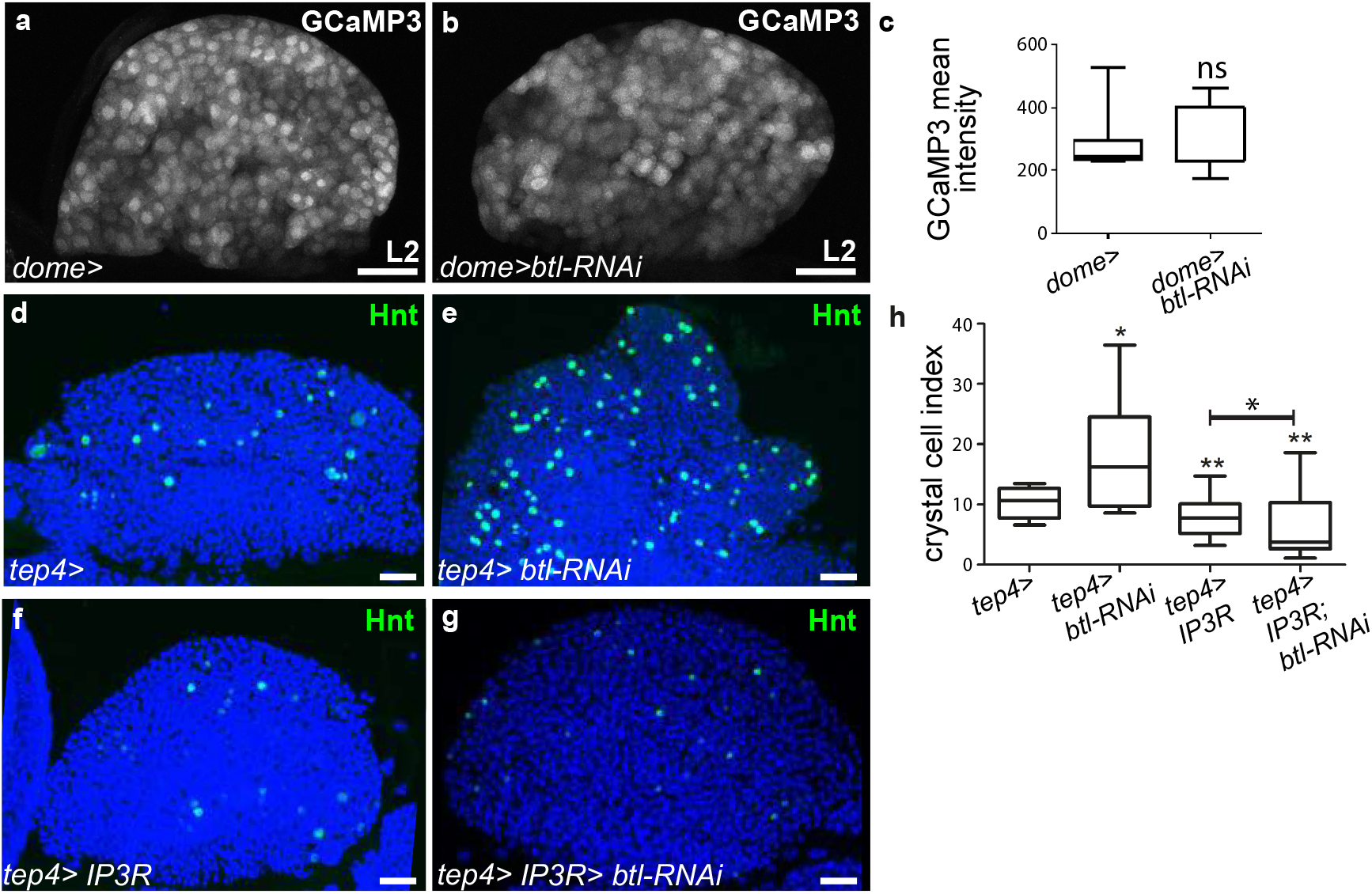
Ca^2+^ levels in progenitors regulate crystal cell differentiation. (a, b) L2 larval lymph glands and the GCaMP3 Ca^2+^ sensor (*dome>UAS-GCaMP3*) is white. No difference observed in GCaMP3 intensity between the control (a) and when *btl is* knocked down in MZ progenitors (b). (c) GCaMP3 intensity. (d-e, f-g) Crystal cells are labelled with Hnt (green) antibody. A decrease in crystal cell numbers is observed when *IP3R* is knocked down in progenitors using the *tep4* driver (f) compared to the control (d). Co-expression of *IP3R* and *btl*-RNAi in progenitors prevents crystal cell differentiation (g) compared to the control (f). (h) Crystal cell index.

## References

Ameri, J., Stahlberg, A., Pedersen, J., Johansson, J.K., Johannesson, M.M., Artner, I., and Semb, H. (2010). FGF2 specifies hESC-derived definitive endoderm into foregut/midgut cell lineages in a concentration-dependent manner. Stem Cells 28, 45–56.

Asada, N., Takeishi, S., and Frenette, P.S. (2017). Complexity of bone marrow hematopoietic stem cell niche. Int J Hematol 106, 45–54.

Baldeosingh, R., Gao, H., Wu, X., and Fossett, N. (2018). Hedgehog signaling from the Posterior Signaling Center maintains U-shaped expression and a prohemocyte population in Drosophila. Dev Biol.

Banerjee, U., Girard, J.R., Goins, L.M., and Spratford, C.M. (2019). Drosophila as a Genetic Model for Hematopoiesis. Genetics 211, 367–417.

Beiman, M., Shilo, B.Z., and Volk, T. (1996). Heartless, a Drosophila FGF receptor homolog, is essential for cell migration and establishment of several mesodermal lineages. Genes Dev 10, 2993–3002.

Benmimoun, B., Polesello, C., Haenlin, M., and Waltzer, L. (2015). The EBF transcription factor Collier directly promotes Drosophila blood cell progenitor maintenance independently of the niche. Proc Natl Acad Sci U S A 112, 9052–9057.

Bischof, J., Maeda, R.K., Hediger, M., Karch, F., and Basler, K. (2007). An optimized transgenesis system for Drosophila using germ-line-specific phiC31 integrases. Proc Natl Acad Sci U S A 104, 3312–3317.

Calvi, L.M., Adams, G.B., Weibrecht, K.W., Weber, J.M., Olson, D.P., Knight, M.C., Martin, R.P., Schipani, E., Divieti, P., Bringhurst, F.R., et al. (2003). Osteoblastic cells regulate the haematopoietic stem cell niche. Nature 425, 841–846.

Calvi, L.M., and Link, D.C. (2015). The hematopoietic stem cell niche in homeostasis and disease. Blood 126, 2443–2451.

Cook, M.S., Cazin, C., Amoyel, M., Yamamoto, S., Bach, E., and Nystul, T. (2017). Neutral Competition for Drosophila Follicle and Cyst Stem Cell Niches Requires Vesicle Trafficking Genes. Genetics 206, 1417–1428.

Crozatier, M., Ubeda, J.M., Vincent, A., and Meister, M. (2004). Cellular immune response to parasitization in Drosophila requires the EBF orthologue collier. PLoS Biol 2, E196.

Dragojlovic-Munther, M., and Martinez-Agosto, J.A. (2013). Extracellular matrix-modulated Heartless signaling in Drosophila blood progenitors regulates their differentiation via a Ras/ETS/FOG pathway and target of rapamycin function. Dev Biol 384, 313–330.

Du, L., Sohr, A., Yan, G., and Roy, S. (2018). Feedback regulation of cytoneme-mediated transport shapes a tissue-specific FGF morphogen gradient. Elife 7.

Dzierzak, E., and Bigas, A. (2018). Blood Development: Hematopoietic Stem Cell Dependence and Independence. Cell Stem Cell 22, 639–651.

Evans, C.J., Hartenstein, V., and Banerjee, U. (2003). Thicker than blood: conserved mechanisms in Drosophila and vertebrate hematopoiesis. Dev Cell 5, 673–690.

Glazer, L., and Shilo, B.Z. (1991). The Drosophila FGF-R homolog is expressed in the embryonic tracheal system and appears to be required for directed tracheal cell extension. Genes Dev 5, 697–705.

Grigorian, M., Liu, T., Banerjee, U., and Hartenstein, V. (2013). The proteoglycan Trol controls the architecture of the extracellular matrix and balances proliferation and differentiation of blood progenitors in the Drosophila lymph gland. Dev Biol 384, 301–312.

Gryzik, T., and Muller, H.A. (2004). FGF8-like1 and FGF8-like2 encode putative ligands of the FGF receptor Htl and are required for mesoderm migration in the Drosophila gastrula. Curr Biol 14, 659–667.

Haas, S., Trumpp, A., and Milsom, M.D. (2018). Causes and Consequences of Hematopoietic Stem Cell Heterogeneity. Cell Stem Cell 22, 627–638.

Hartenstein, V. (2006). Blood cells and blood cell development in the animal kingdom. Annu Rev Cell Dev Biol 22, 677–712.

He, N., Zhang, L., Cui, J., and Li, Z. (2014). Bone marrow vascular niche: home for hematopoietic stem cells. Bone Marrow Res 2014, 128436.

Itkin, T., Kaufmann, K.B., Gur-Cohen, S., Ludin, A., and Lapidot, T. (2013). Fibroblast growth factor signaling promotes physiological bone remodeling and stem cell self-renewal. Curr Opin Hematol 20, 237–244.

Itkin, T., Ludin, A., Gradus, B., Gur-Cohen, S., Kalinkovich, A., Schajnovitz, A., Ovadya, Y., Kollet, O., Canaani, J., Shezen, E., et al. (2012). FGF-2 expands murine hematopoietic stem and progenitor cells via proliferation of stromal cells, c-Kit activation, and CXCL12 down-regulation. Blood 120, 1843–1855.

Iyengar, L., Wang, Q., Rasko, J.E., McAvoy, J.W., and Lovicu, F.J. (2007). Duration of ERK1/2 phosphorylation induced by FGF or ocular media determines lens cell fate. Differentiation 75, 662–668.

Jarecki, J., Johnson, E., and Krasnow, M.A. (1999). Oxygen regulation of airway branching in Drosophila is mediated by branchless FGF. Cell 99, 211–220.

Jung, S.H., Evans, C.J., Uemura, C., and Banerjee, U. (2005). The Drosophila lymph gland as a developmental model of hematopoiesis. Development 132, 2521–2533.

Kadam, S., McMahon, A., Tzou, P., and Stathopoulos, A. (2009). FGF ligands in Drosophila have distinct activities required to support cell migration and differentiation. Development 136, 739–747.

Kiel, M.J., Yilmaz, O.H., Iwashita, T., Terhorst, C., and Morrison, S.J. (2005). SLAM family receptors distinguish hematopoietic stem and progenitor cells and reveal endothelial niches for stem cells. Cell 121, 1109–1121.

Klambt, C., Glazer, L., and Shilo, B.Z. (1992). breathless, a Drosophila FGF receptor homolog, is essential for migration of tracheal and specific midline glial cells. Genes Dev 6, 1668–1678.

Kobayashi, H., Suda, T., and Takubo, K. (2016). How hematopoietic stem/progenitors and their niche sense and respond to infectious stress. Exp Hematol 44, 92–100.

Krzemien, J., Dubois, L., Makki, R., Meister, M., Vincent, A., and Crozatier, M. (2007). Control of blood cell homeostasis in Drosophila larvae by the posterior signalling centre. Nature 446, 325–328.

Lanot, R., Zachary, D., Holder, F., and Meister, M. (2001). Postembryonic hematopoiesis in Drosophila. Dev Biol 230, 243–257.

Lemaitre, B., and Hoffmann, J. (2007). The host defense of Drosophila melanogaster. Annu Rev Immunol 25, 697–743.

Letourneau, M., Lapraz, F., Sharma, A., Vanzo, N., Waltzer, L., and Crozatier, M. (2016). Drosophila hematopoiesis under normal conditions and in response to immune stress. FEBS Lett.

Lin, L., and Affolter, M. (2009). Clonal analysis of growth behaviors druring “Drosophila” larval tracheal development In Doctoral thesis (university of Basel).

Louradour, I., Sharma, A., Morin-Poulard, I., Letourneau, M., Vincent, A., Crozatier, M., and Vanzo, N. (2017). Reactive oxygen species-dependent Toll/NF-kappaB activation in the Drosophila hematopoietic niche confers resistance to wasp parasitism. Elife 6.

Mandal, L., Martinez-Agosto, J.A., Evans, C.J., Hartenstein, V., and Banerjee, U. (2007). A Hedgehog- and Antennapedia-dependent niche maintains Drosophila haematopoietic precursors. Nature 446, 320–324.

McGuire, S.E., Mao, Z., and Davis, R.L. (2004). Spatiotemporal gene expression targeting with the TARGET and gene-switch systems in Drosophila. Sci STKE 2004, pl6.

Monier, B., Astier, M., Semeriva, M., and Perrin, L. (2005). Steroid-dependent modification of Hox function drives myocyte reprogramming in the Drosophila heart. Development 132, 5283–5293.

Morin-Poulard, I., Sharma, A., Louradour, I., Vanzo, N., Vincent, A., and Crozatier, M. (2016). Vascular control of the Drosophila haematopoietic microenvironment by Slit/Robo signalling. Nat Commun 7, 11634.

Morrison, S.J., and Scadden, D.T. (2014). The bone marrow niche for haematopoietic stem cells. Nature 505, 327–334.

Muha, V., and Muller, H.A. (2013). Functions and Mechanisms of Fibroblast Growth Factor (FGF) Signalling in Drosophila melanogaster. Int J Mol Sci 14, 5920–5937.

Nakai, J., Ohkura, M., and Imoto, K. (2001). A high signal-to-noise Ca(2+) probe composed of a single green fluorescent protein. Nat Biotechnol 19, 137–141.

Ornitz, D.M., and Itoh, N. (2015). The Fibroblast Growth Factor signaling pathway. Wiley Interdiscip Rev Dev Biol 4, 215–266.

Oyallon, J., Vanzo, N., Krzemien, J., Morin-Poulard, I., Vincent, A., and Crozatier, M. (2016). Two Independent Functions of Collier/Early B Cell Factor in the Control of Drosophila Blood Cell Homeostasis. PLoS One 11, e0148978.

Pares, G., and Ricardo, S. (2016). FGF control of E-cadherin targeting in the Drosophila midgut impacts on primordial germ cell motility. J Cell Sci 129, 354–366.

Popichenko, D., Sellin, J., Bartkuhn, M., and Paululat, A. (2007). Hand is a direct target of the forkhead transcription factor Biniou during Drosophila visceral mesoderm differentiation. BMC Dev Biol 7, 49.

Rivas, M.L., Cobreros, L., Zeidler, M.P., and Hombria, J.C. (2008). Plasticity of Drosophila Stat DNA binding shows an evolutionary basis for Stat transcription factor preferences. EMBO Rep 9, 1114–1120.

Roy, S., Hsiung, F., and Kornberg, T.B. (2011). Specificity of Drosophila cytonemes for distinct signaling pathways. Science 332, 354–358.

Saito, K., Maeda, M., and Katada, T. (2017). Regulation of the Sar1 GTPase Cycle Is Necessary for Large Cargo Secretion from the Endoplasmic Reticulum. Front Cell Dev Biol 5, 75.

Sato, M., and Kornberg, T.B. (2002). FGF is an essential mitogen and chemoattractant for the air sacs of the drosophila tracheal system. Dev Cell 3, 195–207.

Shim, J., Mukherjee, T., Mondal, B.C., Liu, T., Young, G.C., Wijewarnasuriya, D.P., and Banerjee, U. (2013). Olfactory control of blood progenitor maintenance. Cell 155, 1141–1153.

Sutherland, D., Samakovlis, C., and Krasnow, M.A. (1996). branchless encodes a Drosophila FGF homolog that controls tracheal cell migration and the pattern of branching. Cell 87, 1091–1101.

Thackeray, J.R., Gaines, P.C., Ebert, P., and Carlson, J.R. (1998). small wing encodes a phospholipase C-(gamma) that acts as a negative regulator of R7 development in Drosophila. Development 125, 5033–5042.

Tokusumi, T., Shoue, D.A., Tokusumi, Y., Stoller, J.R., and Schulz, R.A. (2009). New hemocyte-specific enhancer-reporter transgenes for the analysis of hematopoiesis in Drosophila. Genesis 47, 771–774.

Tokusumi, Y., Tokusumi, T., Stoller-Conrad, J., and Schulz, R.A. (2010). Serpent, suppressor of hairless and U-shaped are crucial regulators of hedgehog niche expression and prohemocyte maintenance during Drosophila larval hematopoiesis. Development 137, 3561–3568.

Turner, N., and Grose, R. (2010). Fibroblast growth factor signalling: from development to cancer. Nat Rev Cancer 10, 116–129.

Volk, T., Wang, S., Rotstein, B., and Paululat, A. (2014). Matricellular proteins in development: perspectives from the Drosophila heart. Matrix Biol 37, 162–166.

Yorimitsu, T., Sato, K., and Takeuchi, M. (2014). Molecular mechanisms of Sar/Arf GTPases in vesicular trafficking in yeast and plants. Front Plant Sci 5, 411.

Yu, S., Luo, F., and Jin, L.H. (2018). The Drosophila lymph gland is an ideal model for studying hematopoiesis. Dev Comp Immunol 83, 60–69.

Zhao, J.L., and Baltimore, D. (2015). Regulation of stress-induced hematopoiesis. Curr Opin Hematol 22, 286–292.

Zhao, M., Ross, J.T., Itkin, T., Perry, J.M., Venkatraman, A., Haug, J.S., Hembree, M.J., Deng, C.X., Lapidot, T., He, X.C., et al. (2012). FGF signaling facilitates postinjury recovery of mouse hematopoietic system. Blood 120, 1831–1842.

